# Evolution of neuronal anatomy and circuitry in two highly divergent nematode species

**DOI:** 10.1101/595025

**Authors:** Ray L. Hong, Metta Riebesell, Daniel J. Bumbarger, Steven J. Cook, Heather R. Carstensen, Tahmineh Sarpolaki, Luisa Cochella, Jessica Castrejon, Eduardo Moreno, Bogdan Sieriebriennikov, Oliver Hobert, Ralf J. Sommer

## Abstract

The nematodes *C. elegans* and *P. pacificus* populate diverse habitats and display distinct patterns of behavior. To understand how their nervous systems have diverged, we undertook a detailed examination of the neuroanatomy of the chemosensory system of *P. pacificus*. Using independent features such as cell body position, axon projections and lipophilic dye uptake, we have assigned homologies between the amphid neurons, their first-layer interneurons, and several internal receptor neurons of *P. pacificus* and *C. elegans.* We found that neuronal number and soma position are highly conserved. However, the morphological elaborations of several amphid cilia are different between them, most notably in the absence of ‘winged’ cilia morphology in *P. pacificus*. We established a synaptic wiring diagram of amphid sensory neurons and amphid interneurons in *P. pacificus* and found striking patterns of conservation and divergence in connectivity relative to *C. elegans*, but very little changes in relative neighborhood of neuronal processes.

**Impact Statement:** The substrate for evolutionary divergence does not lie in changes in neuronal cell number or targeting, but rather in sensory perception and synaptic partner choice within invariant, prepatterned neuronal processes.

## Introduction

Comparative studies on nervous system anatomy have a long tradition of offering fundamental insights into the evolution of nervous systems and, consequently, the evolution of behavior (Schmidt-Rhaesa, 2007). Traditionally, such comparative studies have relied on relatively coarse anatomical and morphometric comparisons. The relative simplicity of nematode nervous systems (Schafer, 2016) facilitates the determination and subsequent comparison of neuroanatomical features of distinct nematode species, thereby enabling an understanding of how members of the same phylum, sharing a common body plan, can engage in very distinct behaviors.

We examine here specific neuroanatomical features, from subcellular specializations to synaptic connectivity, of the nematode *Pristionchus pacificus* and compare these features with those of the nematode *Caenorhabditis elegans*. The species shared their last common ancestor around 100 million years ago, which is longer than the human-mouse separation (nei and Glazko, 2001; Prabh et al., 2018; Rota-Stabelli et al., 2013; Werner et al., 2018), and have since diverged to populate very discrete habitats and engage in distinct sets of behaviors. *C. elegans* is a free-living nematode that can mainly be found in rotten fruit while members of the genus *Pristionchus* are regularly found in association with several species of scarab beetles {Herrmann 2006b; Herrmann 2007, (Koneru et al., 2016; Ragsdale, 2015). Species-specific entomophilic beetle association in *Pristionchus* is corroborated by several adaptations. First, *Pristionchus* nematodes exhibit chemosensory responses towards insect pheromones and volatile plant compounds {Hong 2008a; Hong 2008b(Cinkornpumin et al., 2014). Second, *P. pacificus* dauer larvae secrete a high molecular weight wax ester that promotes collective host finding (Penkov et al., 2014). Finally, *Pristionchus* species show predatory behavior towards *C. elegans* and other nematodes (Bento et al., 2010; Liu et al., 2018). All these behavioral features likely require substantial modifications in the nervous system.

One obvious potential substrate for evolutionary adaptions to distinct ecological habitats and interactions with other species is the perception and processing of sensory information. The amphid sensilla, comprised of a pair of bilaterally symmetrical sensilla in the head that are open to the external environment, are the largest nematode chemosensory organs (Bargmann, 2006; Bargmann and Horvitz, 1991; Bargmann et al., 1993). In the model organism *C. elegans*, the anterior sensilla include the amphid sensilla, twelve inner and six outer labial neurons, and four cephalic neurons, along with their associated sheath and socket support cells (Ward et al., 1975; Ware et al., 1975). There are also additional sensory receptors without glia broadly grouped into non-ciliated (URX, URY, URA, URB) and ciliated (BAG, FLP) neurons that are involved in gas sensing, mechanoreception, and male mate-searching behavior (Barrios et al., 2012; Chatzigeorgiou and Schafer, 2011; Doroquez et al., 2014; Gray et al., 2004; Hallem and Sternberg, 2008; Hallem et al., 2011; Perkins et al., 1986). Comparisons of the *C. elegans* amphid neuroanatomy to those of free-living and parasitic nematodes such as *Acrobeles complexus* {Bumbarger 2009}, *Strongyloides stercoralis* {Li 2000; Li 2001}, and *Parastrongyloides trichosuri* {Zhu 2011} have shown that the number and arrangement of the amphid neurons are broadly conserved. However, a more fine-grained comparative analysis of distinct sensory structures as well as their connection to downstream circuits has been largely lacking so far.

Using the 3D reconstructions of serial thin section transmission electron microscopy (TEM), we describe here detailed features of the sensory anatomy of *P. pacificus* as well as the synaptic wiring of sensory neurons to their main, postsynaptic interneurons. Comparing anatomical features of sensory circuitry, from ciliated sensory endings to soma and axon position to synaptic connectivity, we reveal striking patterns of similarities and dissimilarities. The most striking similarity is the overall conservation of neuronal soma and process positioning while the most striking patterns of divergences lie in fine structural details of sensory anatomy as well as synaptic connectivity. These findings demonstrate the existence of several constraints in patterning the nervous system and suggest that major substrates for evolutionary novelty lie in the alterations of dendritic structures and synaptic connectivity.

## Results and Discussion

### Overall Amphid Architecture

Using a combination of 3D reconstructions from TEM sections of two young adult hermaphrodites, as well as live dye uptake and transgene reporter analysis, we set out to characterize the amphid sensory circuitry of *P. pacificus* in order to undertake a comparative analysis with the amphid sensory circuitry of *C. elegans.* For comparison with *C. elegans*, we used electron micographs and findings from both legacy (White et al., 1986) and modern EM methodologies (Doroquez et al., 2014). Despite being approximately 40 years old, the EMs used by John White and colleagues to create The Mind of a Worm remain the most complete publicly available data of the adult hermaphrodite *C. elegans* nervous system. While the methods used to create these legacy EM series are technologically inferior to current practices (chemical fixation, analog microscopy, thicker sections), the overall staining and elucidation of synaptic zones remain useful enough for anatomical comparisons. Recent studies of the amphid dendritic endings in *C. elegans* using the modern High Pressure Freezing (HPF) method have validated the original studies, resulting in a richer description of ultrastructural anatomy (Doroquez et al., 2014). The numerous anatomical similarities and differences we observed across species are both reproducible and share equivalences to previous nematode comparative anatomical studies.

To account for all the amphid neurons in *P. pacificus*, we identified and traced every amphid cilium in the channel from its tip near the mouth to its cell body posterior to the nerve ring in two young adult hermaphrodites. Like *C. elegans*, *P. pacificus* possesses 12 neurons per amphid sensillum. All amphid neurons and the amphid sheath cell in *P. pacificus* have their cell bodies in the lateral ganglion posterior to the nerve ring, which resembles the condition found in *C. elegans* and other nematodes. At the anterior end, the tips of the dendrites are housed by the processes of a pair of amphid sheath cells (AMsh), which are glial cells that connect to the amphidial pore in the cuticle via a pair of amphid socket cells (AMso) that expose the amphid neuronal cilia to the environment (Fig. 1A-C). All amphid neurons have ciliated endings with a circle of varying numbers of doublet microtubules surrounding a few inner singlet microtubules in their transition zones and middle segments (Fig. 1F). We counted 13 cilia (from 11 neurons) in each amphid channel in *P. pacificus,* compared to 10 cilia (from 8 neurons) in the channel in *C. elegans* (Fig. 1D-F, Table 1). The greater number of ciliated endings in *P. pacificus* coincides with a conspicuous lack of neurons with winged morphology found in the *C. elegans* wing cells (AWx), whose elaborate ciliary endings terminate as invaginations inside the distal amphid sheath cell cytoplasm rather than in the amphid channel. This indicates that the AWA, AWB, AWC cellular homologs in *P. pacificus* are among the 11 single or double ciliated neurons in the channel and, therefore, do not display a characteristic “wing”-shaped cilia morphology. Because the wing neurons are the most morphologically distinct and best studied amphid neurons in *C. elegans*, their absence in other nematodes contributes to the difficulty in assigning homology in the amphid neurons in these other nematode species (Ashton et al., 1995; Bumbarger et al., 2007; Bumbarger et al., 2009a; Li et al., 2001; Ward et al., 1975).

**Figure 1.**
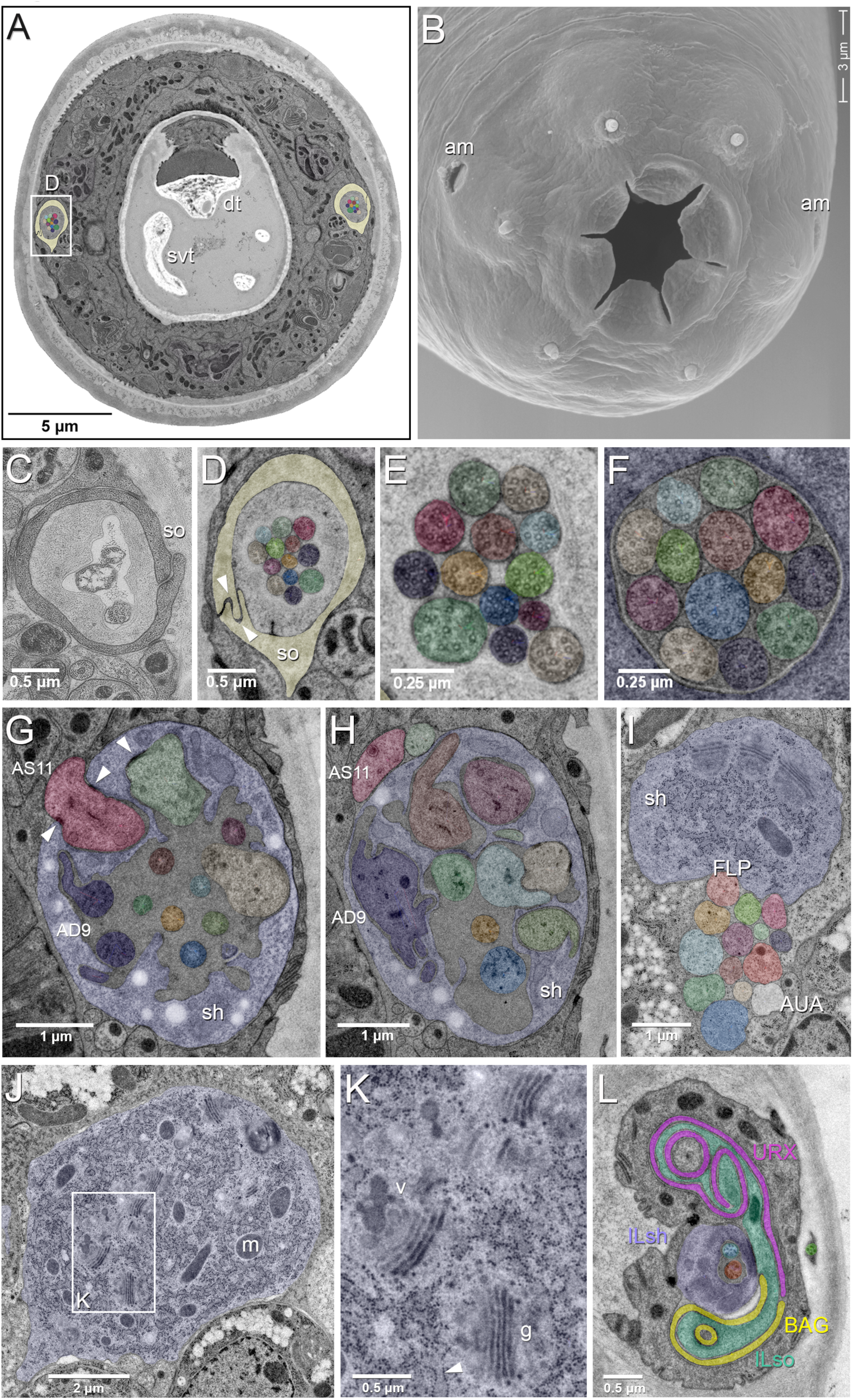
EM of the amphid and two other odor sensing neurons of *P. pacificus* hermaphrodite adults. All images are from specimen 107 (Series 14), except B and C. As specimens were sectioned from the head, left structures appear on the right side in the images and vice versa. (A) Complete transverse section 6.8 µm from the tip of the head showing the amphid socket cells (light yellow) with false-colored amphid cilia bundles inside the channels. Note the dorsal tooth (dt) and sub-ventral tooth (svt) in the buccal cavity. (B) Scanning electron micrograph of the head of an adult animal showing the two amphid openings (am). (C-H) Transverse TEM sections through the amphid cilia at various levels from anterior to posterior (C) Left amphid channel close to the pore, containing the tips of the longest three cilia; the channel is formed by the amphid socket cell (so). (Specimen DB-9-1) (D) Right amphid channel slightly more posterior than in A (7.1 µm from the tip of the head) with all 13 cilia visible in the channel matrix. Arrowheads indicate the autocellular junction of the amphid socket cell (so). (E) Distal segments of the amphid cilia with singlet microtubules in the left channel 6.85 µm from the tip. (F) Middle segments of the right amphid cilia with discernable nine outer doublet microtubules that make up the core axoneme 11.25 µm from the tip; here the channel is formed by the amphid sheath cell (lilac). (G) Section through the region of sheath entry 13.1 µm from the tip, showing the thick periciliary membrane compartment (PCMC) of AS11 (red) entering the left amphid sheath cell (sh), the green and beige PCMCs have just completed their entry. Additionally, the sheath lumen harbors the basal parts of the double cilia of AD9 (dark lilac) with typical transition zone (TZ) arrangement of microtubules and 7 further TZ cilia. Arrowheads mark the adherens junctions between the PCMCs and the amphid sheath. (H) A slightly more posterior section through the left sheath cell (13.85 µm from the tip), showing the distinctively large and irregular outline of the PCMC of AD9 (dark lilac) and the PCMCs of 6 other dendrites. The base of the AS8 cilium (blue) and the TZ of AS5 (orange) are found in the lumen. AS11 is still outside the sheath. (I–K) Transverse TEM sections through the posterior AMsh process and cell body (lilac). (I) Right-side amphid nerve directly anterior of the nerve ring (88.3 µm from the tip) consisting of the AMsh process (sh) with prominent Golgi stacks and the 12 amphid neuron dendrites.. FLP (reddish, just below sheath) and AUA (white) join the amphid process bundle until they diverge from it or end. (J) AMsh cell body (100.8 µm from tip) with numerous mitochondria (m) and Golgi stacks. (K) A higher magnification of the Golgi apparatus (g) and vesicles (v) in the midst of ribosome-studded rough ER cisternae (arrowhead). (L) The bilaterally symmetrical URX and BAG neurons have extended flattened ciliary endings associated with the lateral Inner Labial socket cell (ILso) process; 2.05 μm from the tip. See SI Fig. 6 for the presence of cilia in the URX neurons. Comparable *C. elegans* EM annotations are available at SlidableWorm http://www.wormatlas.org/SW/SW.php/.

**Table 1.** Comparison of amphid neurons in various nematode species. The total number of neurons with dendritic processes encased in the sheath cell in a single amphid compartment is indicated. These neurons are further categorized as neurons with dendrites having single or double ciliated endings in the amphid channel, or as specialized neurons with endings outside the channel but within the sheath cell other than the finger cell.

| Species | Total Neurons | Dendritic ends | Cilia in channel |  |  | Winged cells |
| --- | --- | --- | --- | --- | --- | --- |
|  |  | in channel | Total Count | Single | Double |  |
| *Pristionchus pacificus* | 12 | 11 | 13 | 9 | 2 | 0 |
| *Caenorhabditis elegans*<sup>a</sup> | 12 | 8 | 10 | 6 | 2 | 3 |
| *Haemonchus contortus*<sup>b</sup> | 12 | 10 | 13 | 7 | 3 | 0 |
| *Strongyloides stercoralis*<sup>c</sup> | 13 | 12 | 12 | 12 | 0 | 0 |
| *Acrobeles complexus*<sup>d</sup> | 13 | 12 | 12 | 10 | 1 | 0 |

a. (Ward, 1975)
b. (Li, 2001)
c. (Ashton, 1995)
d. (Bumbarger 2007, 2009)

We designated preliminary names for the *P. pacificus* amphid neurons utilizing a three character nomenclature, in keeping with that used in the descriptions of *C. elegans* (Table 2) (Ward et al., 1975; White et al., 1986). As in *C. elegans,* the first letter A stands for amphid, the second letter is either “S” or “D” for single or double cilia. However, in order to remain initially unbiased, we used numbers rather than letters as the final character. Of the 11 dendrites in the channel nine have single ciliated endings and only two of them, AD9 and AD3, possess double-ciliated endings. The lack of neurons with winged morphology compelled us to consider additional criteria for assigning amphid neuron homology between *P. pacificus* and *C. elegans*, such as the uptake of the live dye DiI, cell body position, and axon projection patterns, which will be described further below.

**Table 2.** Provisional nominations of amphid neuronal homologs between *C. elegans* and *P. pacificus* based on axon projections, live dye DiI uptake, and cell body position. Morphological data based on TEM series of specimen 107 and 148.

| *P. pacificus* neuron | Axon termination site | Entry point into sheath<sup>Δ</sup> | Likely *C. elegans* homolog | Notable features | Cell body Color |
| --- | --- | --- | --- | --- | --- |
| AS1 | dorsal midline | 273/286 dorsal | ASH | Dil uptake | light green |
| AS2 | dorsal midline;<br>not through commissure | 289/304 dorsal | ADL | Dil uptake and DiO; branched axon | violet red |
| AD3 | dorsal midline | 283/308 lateral/<br>dorsal | AWA | *Ppa-odr-3p::rfp* expression | cream |
| AS4 | dorsal midline | 295/313 lateral/ventro-<br>lateral | ASK | Dil uptake;<br>*Ppa-odr-3p::rfp* expression | yellow green |
| AS5 | dorsal midline | 304/320 lateral | ASE | *Ppa-che-1::rfp* expression | orange |
| AS6 | lateral midline | 289/296 lateral/dorsal | ASG | *Ppa-che-1p::rfp* expression;<br>Short axon | light blue |
| AS7 | cross dorsal midline | 298/327 dorsal | AWC | Axon overlap dorsally; *Podr-7p::rfp*; dorsal sheath entry | dark green |
| AS8 | dorsal midline | 313/313 ventral | ASJ | Dil and DiO uptake; ventral sheath entry | middle blue |
| AD9 | dorsal midline | 289/302 dorso-medial | ADF | Dil uptake; *Podr-7p::rfp* shows double cilia and PCMC | dark blue |
| AS10 | dorsal midline | 289/304 dorsal | ASI |  | brown |
| AS11 | dorsal midline | 265/277 dorsal | AWB | Dil uptake | magenta red |
| AF12 | dorsal midline | 576/504 dorsal/<br>ventral | AFD | Finger dendrite morphology | yellow |
| AU1 | dorsal midline | terminates before sheath entry | AUA | Short 'dendritic' process | white |

<sup>A</sup>Based only on Specimen 107: section 0 is the tip of the nose and 3000 is the posterior end of the pharynx, section thickness is about 50 nm (left/right). The first dendrite to enter the amphid sheath is the most posterior (AF12). The dorsal-ventral orientation is with respect to the amphid sheath. Structures of *C. elegans* amphid neurons are available at http://www.wormatlas.org/images/NeuronImageList.jpg.

### The Finger Neuron

The most unambiguous amphid neuron homolog is the finger neuron (which we initially called AF12) with finger-like ciliated endings conserved among *P. pacificus, C. elegans,* and *H. contortus* (Fig. 2A, 2C). The sensory ending of the AF12 dendrite is formed by a short cilium of about 1 µm length and a complex of 30-40 microvilli-like projections branching off from the periciliary membrane compartment (PCMC), which strongly resembles the morphology of the AFD neuron responsible for thermosensation in *C. elegans* and *H. contortus*. We also observed that the *P. pacificus* AF12 sensory ending resides in a more posterior position separate from the other amphid dendrites and has a longer cilium relative to the *C. elegans* AFD (1050 nm compared to 571 nm for *C. elegans* in Fig. 12A)(Doroquez et al., 2014), but nevertheless clearly qualifies as the *P. pacificus* homolog of AFD. The cell body of AF12(AFD) is the most anterior among the amphid neurons, located just ventral of the lateral mid-line. Thus, of the five nematode species with detailed descriptions of their amphidial sensory neuroanatomy, the only nematode species known so far not to share the conservation of finger-like dendritic endings is the mammalian parasite *Strongyloides stercoralis*, which has instead evolved a lamellar morphology for its putative thermosensory neuron ALD (Ashton et al., 1995). Like most other thermosensory neurons, the cilia of the *P. pacificus* finger neurons stay fully embedded inside the amphid sheath cell process, have no contact with the matrix-filled lumen of the sheath cell and do not enter the amphid channel. However, confirmation of the AF12(AFD) neurons’ role in thermosensation will ultimately depend on cell ablation experiments followed by functional behavioral assays.

**Figure 2.**
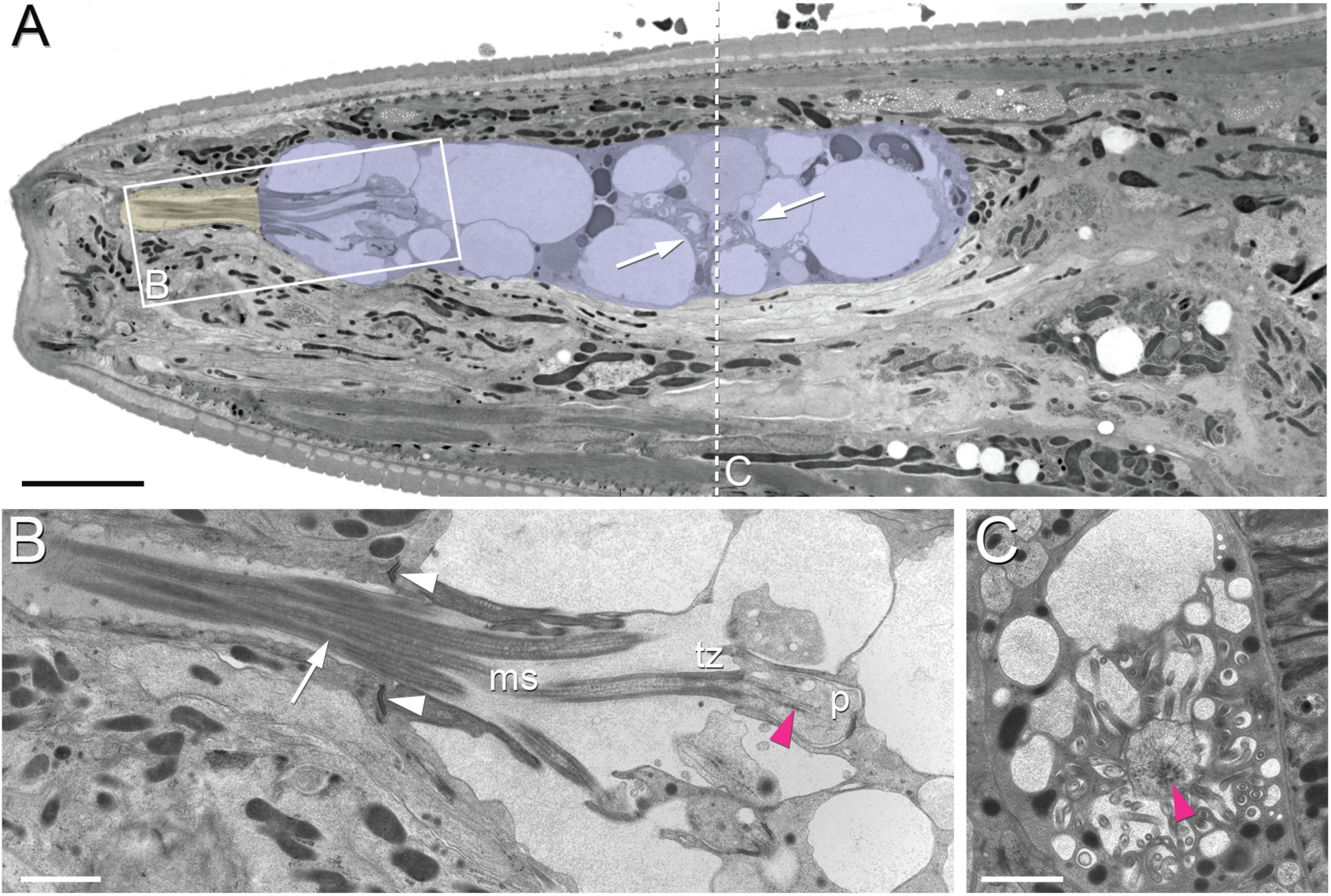
(A) A sagittal TEM section of a *P. pacificus* young adult hermaphrodite. The yellowish shading highlights the narrow amphid channel, which is formed by the socket cell process and contains the distal segments of the amphid cilia. The lilac shading highlights the expanded region of the amphid sheath cell process, which harbors the proximal segments of the amphid cilia in its anterior lumen and, more posteriorly, the finger cell in its cytoplasm amidst a multitude of vesicles. Arrows point at the space taken up by the finger cell cilium with its projections. Scale bar: 5 µm. (B) Detail of A (from a neighboring section) showing the different ciliary regions: middle segment (ms), transition zone (tz), and periciliary membrane compartment (p). The arrow points at cilia in the narrow, matrix-filled amphid channel. Arrowheads mark the adherens junctions between the amphid sheath and socket cell processes. The red arrowhead indicates the cilicary rootlet within the periciliary membrane compartment (p) of one of the amphid neurons. Scale bar: 1 µm. (C) Transverse section through the periciliary membrane compartment of the amphid finger cell with central rootlet (red arrow head) and numerous fingerlike villi in various orientations. The dotted line in A represents the approximate plane of sectioning. Scale bar: 1 µm. Comparative sagittally sectioned *C. elegans* nose images are present at http://www.wormatlas.org/hermaphrodite/neuronalsupport/jump.html?newLink=mainframe.htm&newAnchor=Amphidsensilla31 (Figure 35)

### Cell body positioning and axonal projections of other amphid neurons

Without other signature dendritic endings to nominate possible amphid homologs, we turned to likely conservations in cell body position as well as in the projection trajectories of individual axon processes that enter and terminate in the nerve ring. Using the reconstructions of EM sections (specimen 107, 148; Fig. 6; SI Fig. 5), we looked for *P. pacificus* neurons that share another unique feature of the *C. elegans* AWC neurons: the AWC axons cross the dorsal midline, overlap each other and terminate just before reaching the lateral midline in *C. elegans.* We identified only one pair of single ciliated amphid neurons that shares this unique feature, the AS7 (Table 2). The AS7(AWC) cell body is located between those of the AS1 and AS8 neurons, which are likely the respective cellular homologs of ASH and ASJ based on conservation in DiI uptake and cell body positions along the ventral edge of the amphid neuron cluster (discussed below). Therefore it appears that the homologous AS7 and AWC axons both cross the dorsal midline and terminate above the lateral midline, although the AS7(AWC) cell body appears to have shifted from a position ventral of ASH in *C. elegans* to a position between AS1(ASH) and AS8(ASJ) in *P. pacificus*, such that the three cell types are just ventral and parallel to the lateral midline on each side.

Two other amphid neurons also show strong resemblance to their *C. elegans* counterparts according to their unique axon projections. The *P. pacificus* AS6 is likely the homolog of the *C. elegans* ASG neuron because these counterparts have the unique property of short axon projections that do not project into the nerve ring much further than the lateral midline, terminating before the dorsal midline. The *P. pacificus* AS2 is the likely homolog of the *C. elegans* ADL neuron because like in ADL, the AS2 axons are the only amphid sensory neurons that do not run through the amphidial commissure but enter the nerve ring directly from an anterior projection before branching in the dorsal-ventral direction, which for *C. elegans* ADL, is unique among all amphid neurons (Table 2). If axon projection is more highly conserved than cilia branching, then the *P. pacificus* cellular homolog for ADL is single ciliated and not double ciliated. The nomination of AS2 as the ADL homolog is corroborated by its cell body position just posterior to AS4 and its ability to take up DiI. Altogether, the conserved unique axon trajectories of AS2, AS6, and AS7 argue they are the likely homologs of the *C. elegans* ADL, ASG, and AWC neurons, respectively.

### The presumptive taste receptor neuron pair ASE in *P. pacificus* and *C. elegans*

In *C. elegans,* there is a third amphid neuron pair apart from AS6(ASG) and AS7(AWC) whose axons do not meet and end in a gap junction in the dorsal midline like the rest: the ASE neuron pair, which are the main taste neurons in *C.elegans* (Bargmann and Horvitz, 1991). In *C. elegans,* the axons of ASEL and ASER cross the dorsal midline around the nerve ring until they end ventrally close to the entry point of the contralateral axon (White et al., 1986). In *P. pacificus*, the cell bodies of the presumptive ASE homolog AS5 are located just ventral to the AS2(ADL) neurons at the second-most posterior position of the amphid sensilla [AS8(ASJ) is the most posterior pair]. The conservation of cell body position and the expression of the *Ppa-che-1p::rfp* transgene reporter (see below) provide independent support to nominate the AS5 neurons as the ASE homolog. However, the *P. pacificus* AS5(ASE) axons do not cross each other at the dorsal midline, but rather terminate at the dorsal midline. At their respective termination points *P. pacificus,* ASEL and ASER make a small gap junction with each other, like all but two amphid sensory neuron pairs do (Table 2). This apparent electrical coupling between the *P. pacificus* ASE homologs is notable because, in *C. elegans* no such gap junctions are formed between the ASEs. The consequent lack of electrical coupling between the *C. elegans* ASEL and ASER neurons has been found to be necessary to produce a physiological asymmetry between these neurons, such that both neurons are differentially activated by distinct sensory cues, i.e. their function is “lateralized” (“left/right asymmetric”)(Ortiz et al., 2009; Suzuki et al.). Moreover, it has been shown that artificial establishment of gap junctions between the left and right *C. elegans* ASEs leads to loss of lateralization and changes in chemotaxis behavior (Rabinowitch et al., 2014). The apparent coupling of ASEL and ASER in *P. pacificus* suggests that these two neurons are not functionally lateralized in *P. pacificus*.

Two additional genetic observations are consistent with a lack of functional lateralization in *P. pacificus*: First, the key regulatory factor that triggers the asymmetry of the ASEL/R neurons in *C. elegans,* the miRNA *lsy-6* (Cochella and Hobert, 2012; Johnston and Hobert, 2003) does not exist in the *P. pacificus* genome (*lsy-6* is specific to the *Caenorhabditis* crown clade)(Ahmed et al., 2013). Second, the ASEL and ASER neurons in *C. elegans* (as well as closely related *Caenorhabditis* species) each express a different subfamily of duplicated and chromosomally-linked, receptor-type guanylyl cyclases (rGCYs), the ASER-rGCYs (e.g. *gcy-1, gcy-2, gcy-3, gcy-4, gcy-5*) and the ASEL-rGCYs (*gcy-6, gcy-7, gcy-14, gcy-20*) (Ortiz et al., 2006; Yu et al., 1997), several members of which are thought to be salt chemoreceptors (Ortiz et al., 2009). In contrast, the *P. pacificus* genome contains no *C. elegans-*ASEL-type rGCYs and only a single ASER-type rGCYs (SI Fig. 1). Together with the changes in electrical coupling of the ASE neurons in *C. elegans* versus *P. pacificus,* the differences in the existence of molecular regulators (*lsy-6*) and molecular effectors (*gcy* genes) of *C. elegans* ASE laterality suggest that the ASE neurons of *P. pacificus* may not be functionally lateralized.

### Patterns of neuronal dye-filling are largely conserved

To further explore homologous features of *P. pacificus* and *C. elegans* amphid neurons, we visualized the organization and location of the neuronal cell bodies and their dendritic processes in live wild-type animals with the lypophilic dyes DiI and DiO. In *C. elegans* non-dauer hermaphrodites, FITC and DiI specifically and routinely stain five pairs of head amphid neurons with open sensory endings to the environment - ASK, ADL, ASI, ASH, ASJ (Fig. 3) - along with two pairs of tail phasmid sensory neurons - PHA and PHB. Additionally, ADF is stained weakly by FITC and AWB by DiI. (Hedgecock et al., 1985; Perkins et al., 1986; Starich et al., 1995). It is unclear why only certain amphid neurons exposed to the environment take up DiI and DiO. In *C. elegans,* the cell bodies of the three dorsal-most amphid neurons just below the amphid sheath cell form a distinctive DiI-stained trio (ASK, ADL, ASI), while ASH and ASJ are visible at different focal planes ventral or posterior to this trio, respectively. The cell ablations of the DiI positive ASH neurons show that its polymodal function is strongly conserved across various nematode species (Srinivasan et al., 2008). While ciliated channel neurons take up DiI differentially but are superficially conserved in diverse free-living nematode species examined, including other *Caenorhabditis* species, *Panagrellus redivivus*, and *P. pacificus*, DiI uptake patterns are less similar in the insect parasites such as *Steinernema carpocapse* and *Heterorhabditis bacteriophora.* Thus, DiI uptake is an important but not singular criterion for defining homologous sensory neurons (Han et al., 2016; Srinivasan et al., 2008).

**Figure 3.**
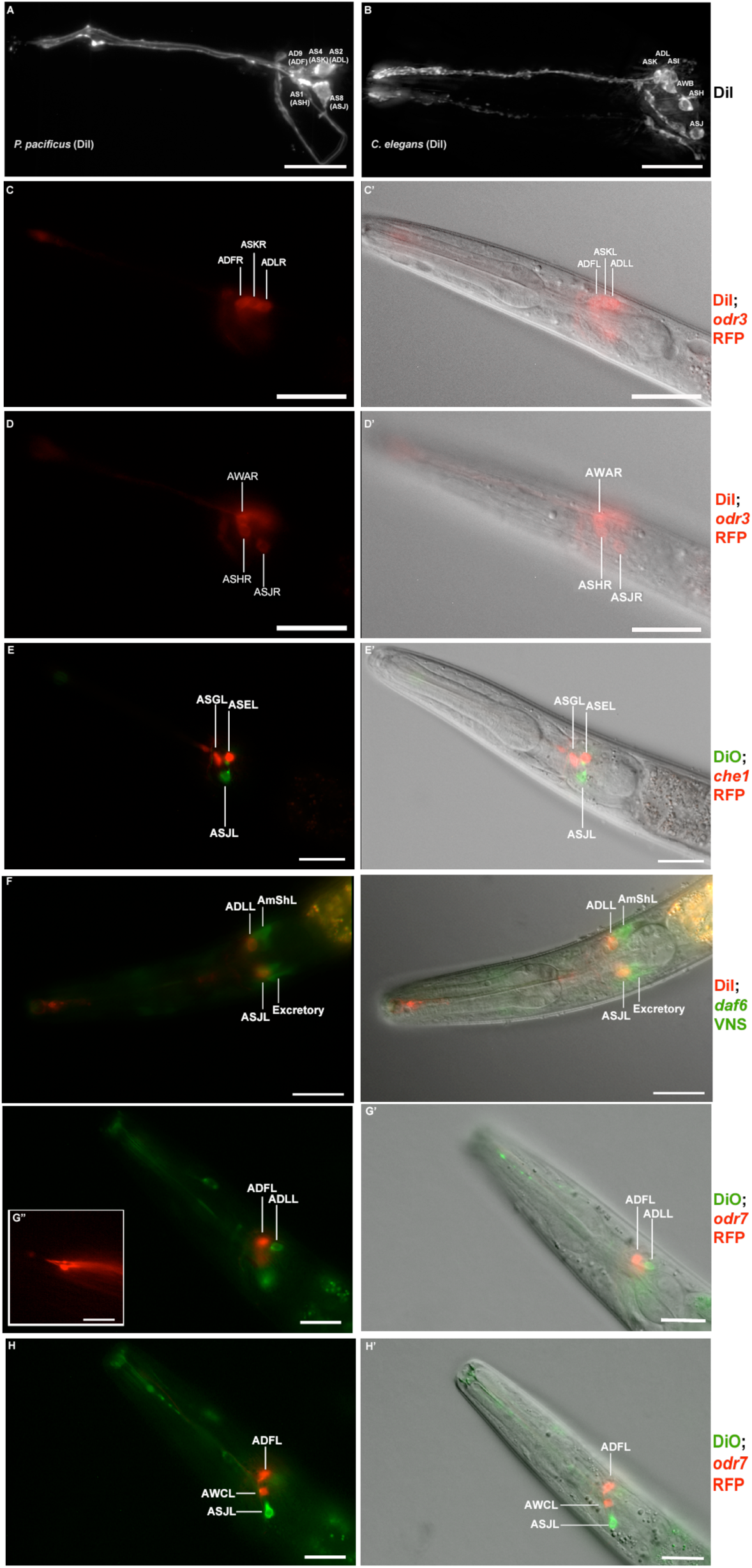
Dye filling and reporter gene expression in *P. pacificus* amphid neurons. (A) Stacked and deconvoluted fluorescent images of DiI stained amphid neurons in a *P. pacificus* young adult hermaphrodite (left lateral view). AD9(ADF), AS4(ASK), AS2(ADL), AS1(ASH), and AS8(ASJ) amphid neurons stain robustly. (B) In *C. elegans,* DiI stains the ASK, ADL, ASI, AWB, ASH, and ASJ neurons. (C-H) Single plane fluorescent images in *P. pacificus*. (C-D) DiI stained *Ppa-odr-3p::rfp* transgenic J3 larvae showing the three dorsal amphid neurons and the larger ASH and ASJ neurons in two focal planes. (E) *Ppa-che-1p::rfp* transgenic J4 hermaphrodite showing expression in ASE and ASG. (F) A DiI stained *Ppa-daf-6p::venus* J4 larva showing ADL and ASJ staining, anterior to the amphid sheath and the excretory cells, respectively. (G, H) A DiO stained *Ppa-odr-7p::rfp* young adult in two focal planes showing dye filling in ADL and ASJ, and RFP expression in ADF and ASK. (G” inset) The dendritic ends of another *Ppa-odr-7p::rfp* adult show the prominent PCMC of ADF with double cilia (bottom), and the smaller PCMC of ASK with a single cilium. Scale bar: 5 µm. (C’-G’) DIC overlay of the same DiI fluorescence images. Anterior is left and dorsal is up. Scale bar: 20 µm. (Representative images based on: *Ppa-odr-3p::rfp* n=22; *Ppa-che-1p::rfp* n=31; *Ppa-daf-6p::venus* n=60; PS312 n=10; *Ppa-odr-7p::venus* n=26). Additional Z-stacks: SI Fig. 8 and 10.

In *P. pacificus* hermaphrodites, DiI stains five pairs of amphid neurons in a pattern similar to those stained in *C. elegans* [AS1(ASH), AS2(ADL), AS4(ASK), AS8(ASJ), and AD9(ADF)](Fig. 3A-B) but DiO stains only the AS2(ADL) and AS8(ASJ) neurons (Fig. 3G-H). In contrast to *C. elegans*, the *P. pacificus* phasmid neurons only dye fill in the dauer larvae (data not shown), while the AS11(AWB) neurons rarely dye fill in any developmental stage with either dye (AWB is stained more robustly with DiO than with DiI in *C. elegans*). The similarity in cell body positions for AS8 and morphology for both AS2 as well as AS8, suggest they are likely ADL and ASJ homologs, respectively. Thus, DiI staining in the three dorsal-most cells in the same focal plane as the AMsh cell body has superficial resemblance to the three dorsal-most neurons in *C. elegans* (Fig. 3; SI Fig. 2). If the posterior AS2 cell in the trio is the ADL homolog, rather than the ASI, then the middle cell of this trio, AS4, is likely the ASK homolog, and the anterior cell of this trio, AD9, is the ADF. Thus AD9(ADF), AS4(ASK), and AS2(ADL) together make up the dorsal trio of neurons that take up DiI. In this proposed homology, the putative ASI homolog, AS10, does not take up DiI. In *C. elegans,* the ASI neurons are important for regulating dauer development and are remodeled in the dauer larvae such that the dauer ASI neurons no longer take up DiI due to cilia retraction from the amphid pore (Albert and Riddle, 1983; Peckol et al., 2001). Given that in *P. pacificus* the phasmid neurons only take up DiI as dauer larvae, it is possible that differential DiI uptake in homologous neurons between the two species recapitulate certain remodeling events in amphid neurons during dauer entry and exit that is lost in one lineage. While the positioning of the neurons taking up DiI appears mostly unaltered at first glance, cell body positions or the chemical environment for the live dye may have diverged modestly during evolution.

### The Amphid Sheath Glia

To complement the characterization of the neuronal composition of the amphid sensillum, we set out to visualize the morphology of the glia-like amphid sheath cells (AMsh cell) in more detail and *in vivo*. To this end, we constructed an AMsh reporter using a 2.4 kb region of the *P. pacificus daf-6* promoter to drive the Red Fluorescent Protein (RFP). DAF-6 is a Patch-related protein required for proper tubule formation in *C. elegans,* including the morphogenesis of the amphid sheath channel (Oikonomou et al., 2011). Indeed, *P. pacificus daf-6* expression shows strong conservation not only in the AMsh, but also in cells of the excretory duct and pore, seam cells, as well as in the VulE epidermal cell that contributes vulva formation during the mid-J4 larval stage (Perens and Shaham, 2005) (Fig. 4A; SI Fig. 3; data not shown). We found a similar *daf-6* expression profile in dauer larvae (Fig. 4B). The cell bodies of the amphid sheath cells are distinctively large and sit dorso-anterior to the terminal bulb of the pharynx. The AMsh cells extend thick processes anteriorly that swell to form paddle- or bottle-like endings near the nose of the worm (Fig 2A,B, Fig. 6A,B), unlike their *C. elegans* counterparts, whose endings fan out widely into the head, forming large sheets to accommodate the expanded ciliated endings of the winged AWC neurons and the finger cells. The amphid socket cells distally surround the ciliated ends of the sensory dendrites, each forming an autocellular junction onto itself to create a pore in the cuticle of the lateral lip that is in direct contact with the environment (Fig. 1C-D; Fig 2A,B). The sensory dendrites enter the matrix-filled lumen of the amphid sheath cell process quite distal to their respective cell bodies. The lumen, or channel, is an extracellular space formed by the merging of numerous large matrix-filled vesicles (Fig. 1G-H, SI Fig. 4). Accordingly, the cytoplasm of the posterior process and the cell body is rich in mitochondria, ribosomes, rough endoplasmic reticulum, Golgi complexes and vesicles of different sizes, distinguishing the AMsh as an active secretory cell (Fig. 1I-K). We speculate that some of the secreted factors act as accessory to chemosensory function to modify odors or water soluble molecules (Bacaj et al., 2008; Cinkornpumin et al., 2014), or to protect the neurons against reactive oxygen species (Liu et al., 2017; Liu et al., 2015).

**Figure 4.**
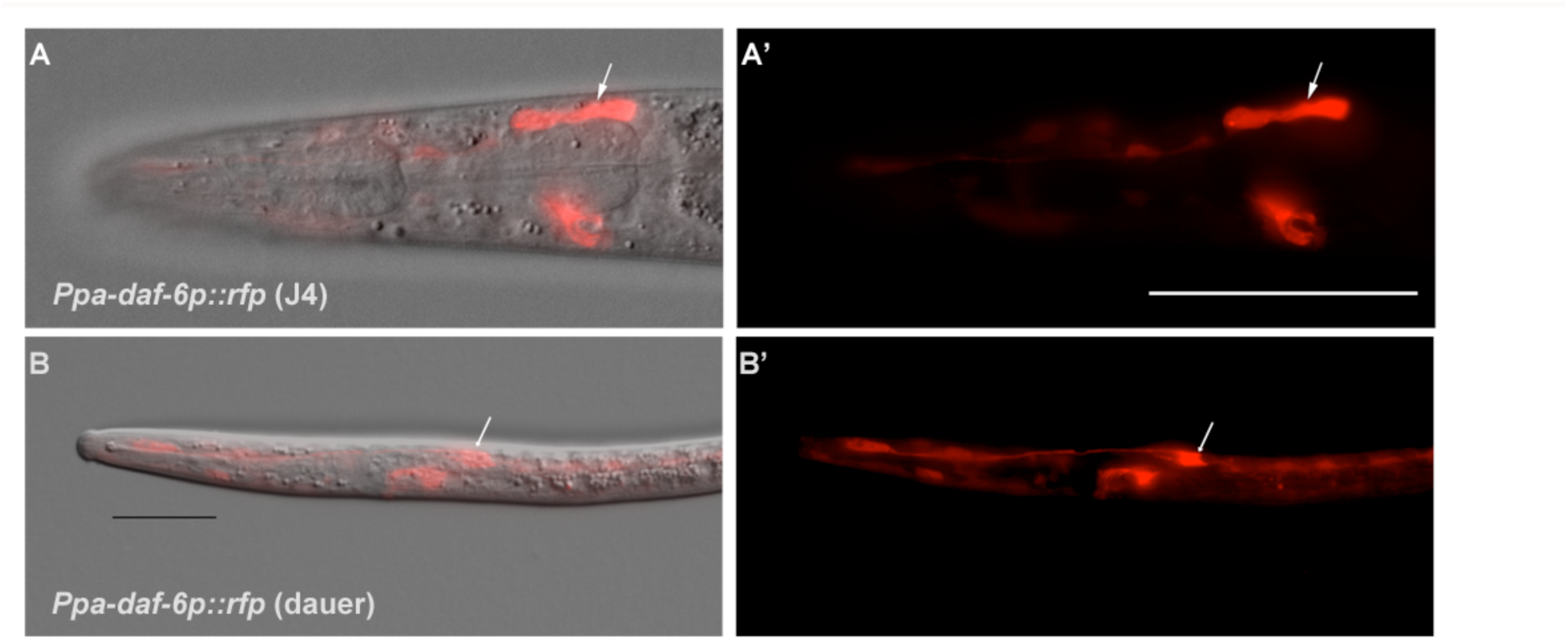
(A-A’) DIC overlay and *Ppa-daf-6p::rfp* expression in the dorsally located amphid sheath cell as well as the excretory cell on the ventral side of a J4 hermaphrodite. Scale bar: 50 µm. (B-B’) DIC overlay and *Ppa-daf-6p::rfp* expression in a dauer larva with prominent amphid sheath, excretory cell, and seam cells expression. Scale bar: 25 µm. Arrows indicate the amphid sheath cell body.

In contrast to *C. elegans*, all the non-finger amphid neurons have dendritic processes that terminate in the amphid channel and are thus in direct contact to the external environment (Fig. 1 A-F, Fig. 2A-B). Because the *C. elegans* AMsh cells accommodate the prominently large AWC winged neurons, the left and the right AMsh can fuse to each other in the dauer larvae, which undergo radial constriction during dauer entry (Procko et al., 2011). Given the absence of any amphid neurons with winged morphology in *P. pacificus*, not surprisingly, *Ppa-daf-6p::rfp* expression in the pair of anterior AMsh endings remains distinctively separated in *P. pacificus* dauer larvae (Fig. 4B). While the *C. elegans* amphid sheath processes swell at their anterior ends to accommodate the wing-shaped cilia, the amphid sheath processes in *P. pacificus* maintain a tube-like form, becoming slightly wider towards the anterior tip, where it takes up the 12 neurons, and thinner posteriorly near the nerve ring in an apparent kink anterior to the cell body (n=3). As a result, the amphid sheath morphology is more tubular than the *C. elegans* amphid sheath.

### Dendrite Entry into the Sheath Cell

Next, we examined the interaction of the neuronal dendrites with sheath cells in more detail. During *C. elegans* development, the nascent dendrites of the amphid neurons and the processes of the sheath glial cells attach to the tip of the nose and from there elongate posteriorly by retrograde growth when the cell bodies start migrating posteriorly to their final position. This elongation process happens in a precisely coordinated way in each bundle (Heiman and Shaham, 2009). Reconstruction of transverse TEM sections from four *C. elegans* specimens show invariant arrangement of the amphid neurons in the sensory channel, such that each amphid neuron can be reliably identified by its position in the channel (Ward et al., 1975). In our data set of only two specimens, we saw a variation in the position of one cilium even between the left and right side in one animal. To determine the degree in which the order and position of each dendrite entry is stereotypical in *P. pacificus*, we determined the individual entry points of amphid dendrites into the sheath in the reference sample (Specimen 107, similar data from specimen 148 not shown; Table 2 and SI Fig. 4). Over most of their length the dendrite bundles are located on the ventral side of the AMsh but from 36.4 µm and 39.5 µm from the tip of the head on the left and right sides, respectively, shortly before the tips of the AMsh processes start to become wider, the bundles loosen and dendrites distribute around the sheath processes, finally arranging themselves into a dorsal and a ventral group. The first dendrites to enter are those of the finger neurons AF12(AFD) at approximately section 25 µm from the tip of the head, or 1/6 the remaining distance between the posterior end of the pharynx (section 3000) and the tip of the ‘nose’ at the anterior end (Section 0). Anterior to the region occupied by AF12(AFD), the dendritic bundle splits into two or three groups consisting of one or more dendrites that enter the sheath in variable order from three sides. For example, AS4 enters laterally together with AD3(AWA), AS5(ASE), AS6(ASG) in the left amphid bundle, or ventrally accompanied by AS5(ASE) and AS8(ASJ) in the right amphid bundle. AS8(ASJ) however, enters ventrally together with AS5(ASE) on both sides after the dendrite bundles split. The entry of the finger neuron can be dorsal or ventral, and the entry of AD3(AWA) and AS6(ASG) can also be lateral or dorsal. In contrast, the site of entry for eight other neurons is consistently dorsal on both sides. The dendrites of AS11(AWB) are the last to enter the sheath (dorsally) 13.5 and 13.8 µm from the tip of the head. Thus, unlike in *C. elegans,* the sequence and site of entry for many neurons relative to the amphid sheath are variable in *P. pacificus*, and additional samples may reveal even more variations in a single specimen.

### Homologies of other sensory neurons

In addition to the sensory sensilla, *P. pacificus* also possesses five types of sensory receptors that terminate in the nose region of the animal but are not accompanied by sheath or socket cells. Of these, the BAG neurons (named for bag-like sensory ending) stand out for being required for sensing carbon dioxide in *C. elegans,* which is presumably also important for host detection in parasitic nematodes (Hallem and Sternberg, 2008; Hallem et al., 2011). In *P. pacificus,* the two BAG neuron cell bodies are located just anterior of the nerve ring in subventral position close to the isthmus (Fig. 5A-B, Fig. 6A, SI Fig. 5), with their dendritic processes running in the ventral-most position of the amphid process and lateral labial bundle. The ciliated distal ends of these processes form branched lamellae around the ventral halves of the lateral inner labial socket cells (ILsoL), opposite to those of another pair of internal receptors, the URX neurons (Fig. 1L, Fig. 5A-B). In *C. elegans,* the pair of URX neurons is known to be important for sensing oxygen levels and to control carbon dioxide response (Carrillo et al., 2013; Gray et al., 2004). The cell body of the URX neuron in both nematode species is located posterior to the nerve ring, and directly anterior to the anterior-most amphid neuron cell body in dorsal position (ADF in *P. pacificus* and ASK in *C. elegans*) (Fig. 6B). Interestingly, the dendritic endings of the homologous *P. pacificus* URX neurons appear to be ciliated with eight microtubules and are associated with the dorsal half of the lateral IL socket cells (Fig. 1L, Fig. 5A, SI Fig. 6), whereas the *C. elegans* URX neurons are non-ciliated and unaffiliated with any sensilla (Doroquez et al., 2014). Ciliated URX neurons are also found in other nematode species, e.g. one of two dendritic endings of the URX neurons in the soil-dwelling nematode *Acrobeles complexus*, as well as all of the URX endings of the mycophagus nematode *Aphelenchus avenae* (Bumbarger et al., 2009b; Ragsdale et al., 2009). Lastly, the four putative URY neurons have dendritic endings with numerous (6-20) singlet microtubules, which split into several overlapping branches extending membranous elaborations towards all six ILsh and ILso, similar to their *C. elegans* counterparts (Doroquez et al., 2014).

**Figure 5.**
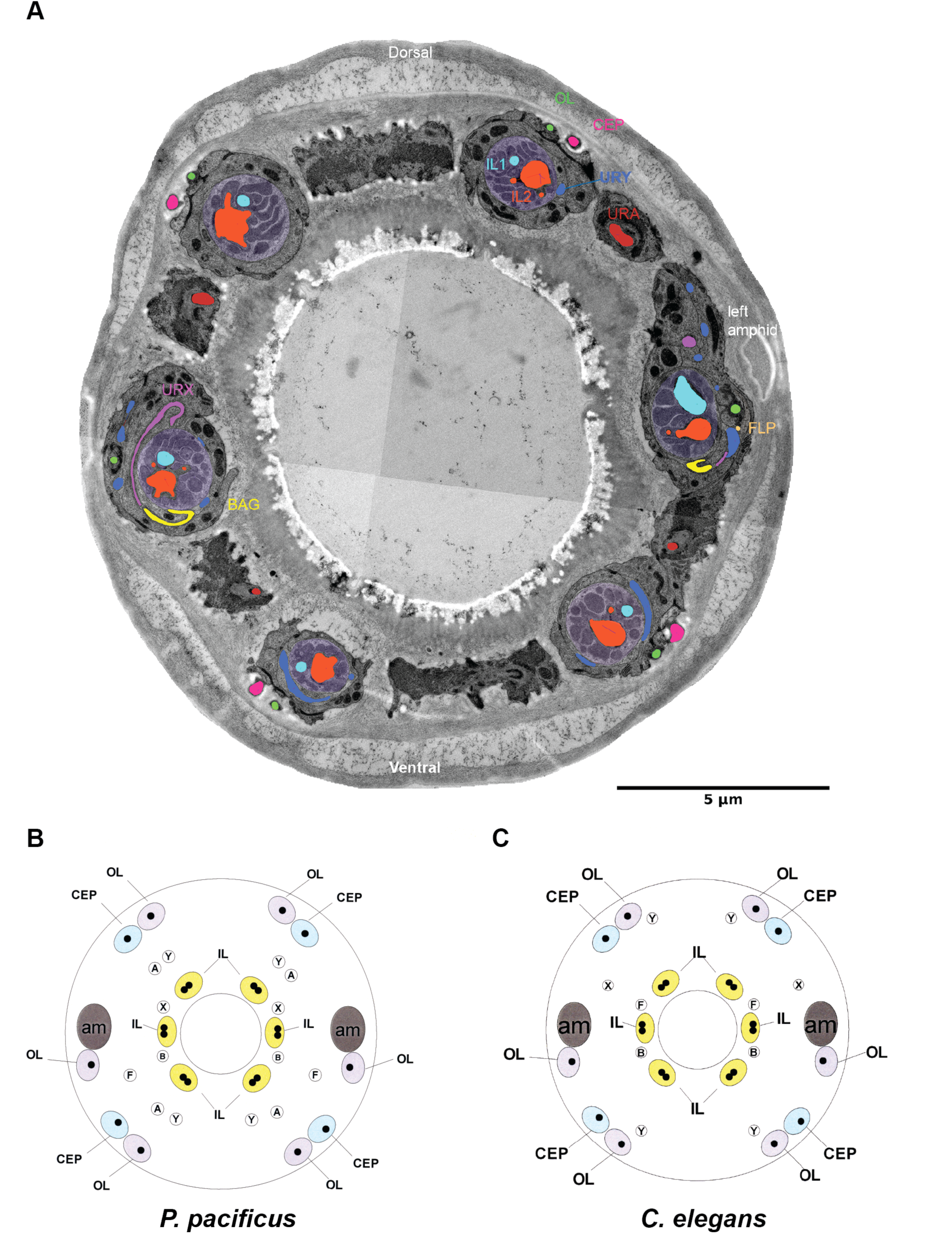
(A) Transversal TEM section of *P. pacificus* cuticular sensilla and internal receptors of the mouth region. One from each sensillum type is labeled in the same false color as the neuron. (B-C) Schematic illustration of the cuticular sensilla in *P. pacificus* and *C. elegans*. Inner Labial 1 and 2 (IL), Outer Labial (OL), Cephalic (CEP), URA (A), URY (Y), URX (X), BAG (B), FLP (F), amphid (am).

**Figure 6.**
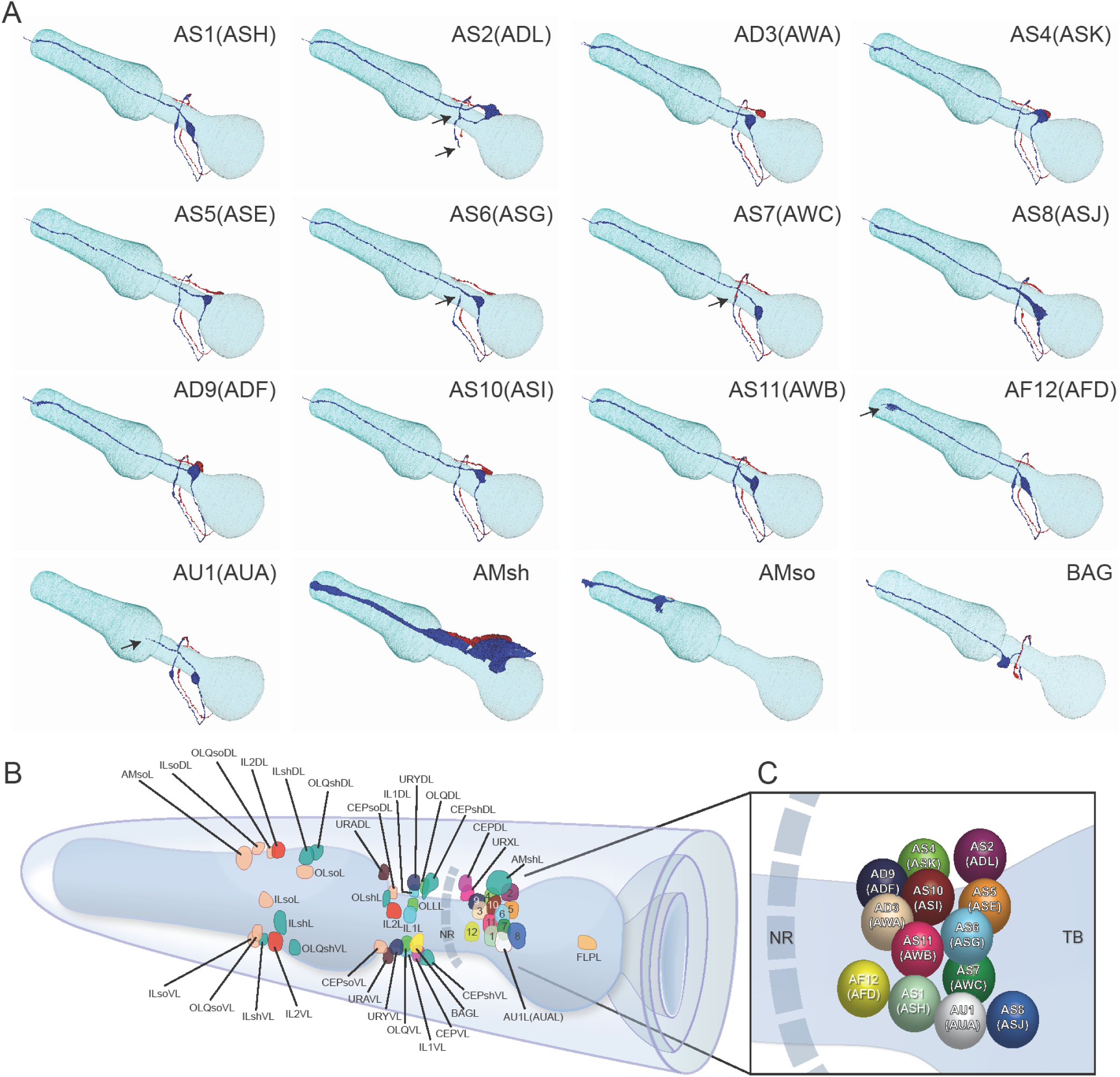
(A) Three-dimensional renderings of individual pairs of amphid neurons, amphid sheath, amphid socket, internal sensory receptor AU1 and BAG neuron in a *P. pacificus* young adult hermaphrodite (Specimen 107). Left lateral view, anterior is to the left. Blue and red renderings denote left and right counterparts, respectively. The pharynx is outlined in teal. Nomenclature is displayed with *P. pacificus* first and *C. elegans* name in parenthesis. Black arrows denote lateral nerve ring entry (not through amphid commissure) and branched axons in AS2 (ASG), short axons ending at the lateral midline in AS6 (ASG), dorsal overlap of axon endings in AS7 (AWC), fingerlike ending of AF12 (AFD) dendrite terminating within the sheath cell posterior to the nose, and short dendritic-like endings of AUA anterior to the nerve ring. Movies with original files can be accessed at SI Fig. 5. (B) Schematic diagram of the positions of the nuclei of the left cuticular sensilla with sheath and socket cells and of the internal receptors. The nerve ring (NR) is indicated by a dotted line. Amphid cell bodies are labelled 1–12 by their last number, with color code the same as in Fig. 1. (C) Schematic of amphid nuclei as seen from the left side with full names, nuclei slightly enlarged. (TB = terminal bulb) Cartoons of *C. elegans* amphid neuron anatomy are present at http://www.wormatlas.org/images/NeuronImageList.jpg. Cartoons of *C. elegans* ganglia are present at http://www.wormatlas.org/images/VCMNganglia.jpg.

We also observed that unlike the FLP neurons in *C. elegans*, the *P. pacificus* FLP neurons do not appear to be associated with the lateral IL socket cell (Doroquez et al., 2014), such that the most anterior process we traced on the left side of Specimen 107 terminates in close proximity to URX and BAG in a region where the BAG neurons start to form their lamellae around the IL socket (Fig. 5A-B). The *C. elegans* FLP neurons have elaborate multi-dendritic structures throughout the head region, and are polymodal receptors for thermosensation and mechanosensation (Albeg et al., 2011; Chatzigeorgiou and Schafer, 2011). Like in *C. elegans*, *P. pacificus* FLP cell bodies are located posterior to all of the amphid neuron cell bodies near the terminal pharyngeal bulb, and have dendrites with extensive branching, whereas their axon projections do not enter the nerve ring. In *C. elegans,* the BAG and FLP neurons are unusual among the anterior sensory neurons for not possessing their own set of glial cells, but instead associate with the socket cells of the lateral IL neurons (Doroquez et al., 2014). Since it is not clear if the *C. elegans* BAG and FLP association with the IL socket cells is primarily for structural stability, i.e. the socket cells could interact with any nearby neuron or provide functional support, it also remains to be determined what functional significance is there, if any, for *P. pacificus* URX neurons rather than the FLP neurons to associate with the lateral IL sensilla.

Another unusual neuron pair, called AUA in *C elegans* (for Amphid Unknown Type A), has a clear homolog in *P. pacificus*. Although not a true amphid sensory neuron, the *C. elegans* AUA also sends its axon through the amphid commissure and possesses a distinctive dendrite-like process (White et al., 1986). While the axons of the AUA neuron pair are similar to many other amphid sensory neurons, also terminating by making a gap junction between the left and right partners at the dorsal midline, their dendrites terminate just anterior of the nerve ring rather than projecting to the anterior end of the animal. We observed a similar neuron in *P. pacificus,* AU1 (Amphid Unknown 1), whose dendritic-like process terminates just anterior of the nerve ring. Given this strong dendritic structural similarity, as well as a similar axonal projection, AU1 is the likely homolog AUA. The conservation of the AU1(AUA) neurons, which mediate oxygen sensing and social feeding behavior in *C. elegans*, strongly implies that they share conserved functions in several nematode species (Bumbarger et al., 2009a; Chang et al., 2006; Coates and de Bono, 2002).

### Axonal process neighborhood is highly conserved

We next explored amphid neuron axon process placement within the nerve ring. The original characterization of the *C. elegans* nervous system identified and used reproducible placement of axons within ganglia as criteria to identify individual neurons (Ware et al., 1975; White et al., 1986)(Fig. 7A). We used a similar approach and compared axonal placement in *P. pacificus* to *C. elegans.* Specifically, we evaluated the ultrastructure of the ventral ganglion (Fig. 7B and 7D), which is proximal to where the amphid commissure enters the main axonal neuropil, as well as the dorsal midline (Fig. 7C and 7E), where most amphid axons terminate. Using the color codes from Table 2, we labeled the location of each amphid axon in both species. Despite differences in sample preparation (HPF for *P. pacificus* and chemical fixation for *C. elegans*) and EM section thickness (50nm for *P. pacificus* and ~80nm for *C. elegans)*, we observed striking similarities in axonal placement in both species, providing further strong support for our homology assignment of neurons. For example, we observed similar amphid sensory neuron axonal fasciculation along the ventrodorsal and mediolateral axes of the ventral ganglion, where the ASJ and ASG neurons are most lateral, and the AFD and AUA neurons are the most ventral in both species (Fig. 7B and 7D). The most posterior segment of the dorsal nerve cord, where most bilaterally symmetric neurons meet, is occupied by a similar set of neurons in both species (Fig. 7C and 7E). Also, of note are the ASJ neurons which are most ventral and ADL which are most lateral in both species.

**Figure 7.**
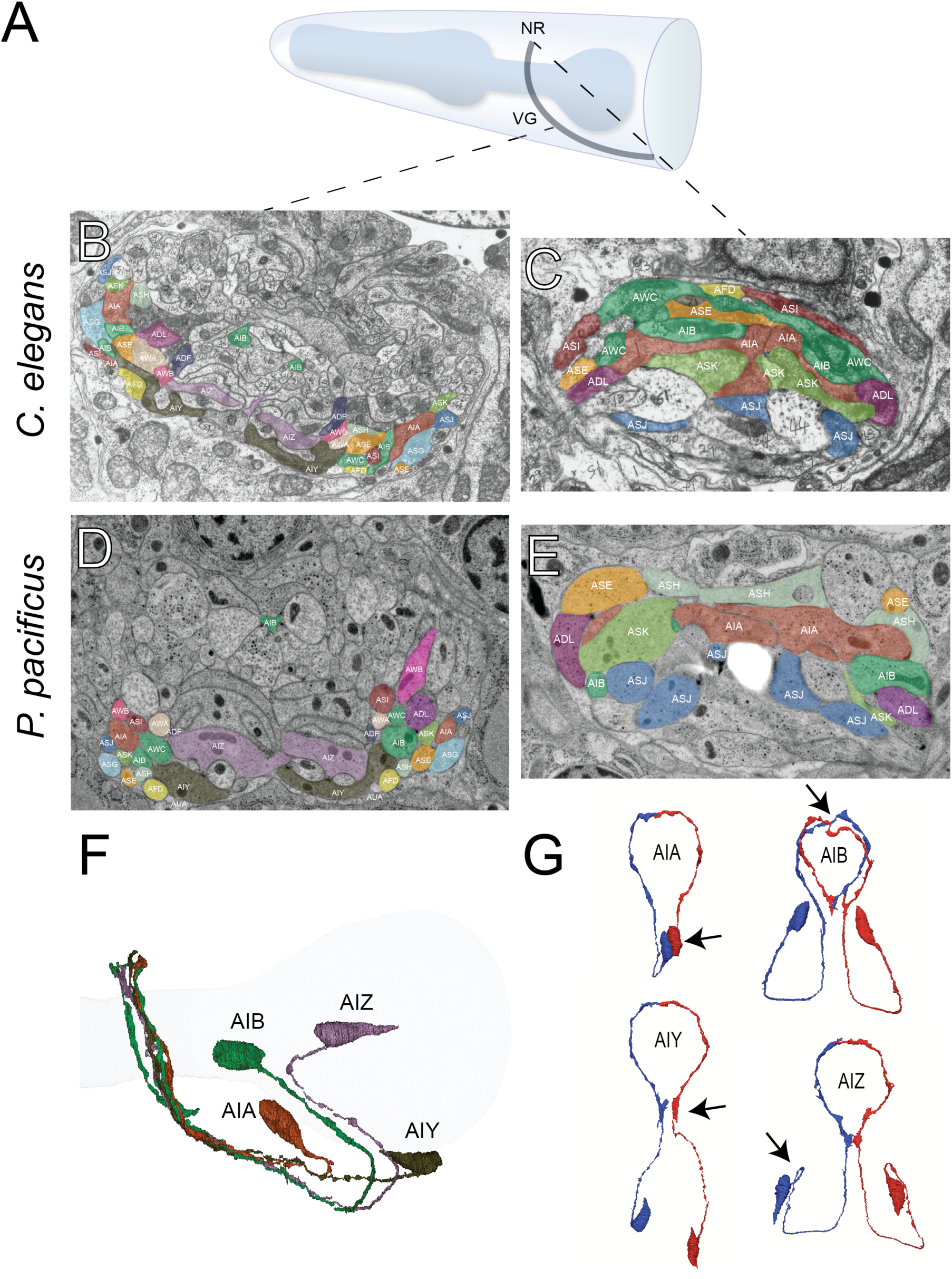
(A) Model of the anterior *P. pacificus* highlighting the nerve ring (NR) and ventral ganglion (VG). (B) Electron micrograph of the anterior *C. elegans* (N2U sample) ventral ganglion and dorsal nerve ring (C) colored with the same color code as in Fig 1. Also shown are AIA (maroon), AIB (dark green), AIY (green/gray), and AIZ (purple). *C. elegans* neuronal nomenclature is used. Electron micrograph of the anterior *P. pacificus* (specimen 107) ventral ganglion (D) and dorsal nerve ring (E) colored with the same color code as in Fig 1. (F) Three dimensional rendering of the AIA, AIB, AIY, and AIZ interneurons from *P. pacificus* (specimen 107) with pharyngeal outline shown in light teal. (G) Individual three dimensional renderings of *P. pacificus* (specimen 107) amphid interneurons shown with a dorsoposterior view with the left and right neurons in blue and red, respectively. Arrows show anatomical features present in both species: AIA cell bodies are adjacent, AIB axons undergo a dorsal neighborhood change within in the nerve ring, AIY neurons have varicosities relating to synaptic output posterior to the nerve ring, and AIZ axons show anterior projections before commissural entry. Amphid neuroanatomy can be compared to *C. elegans* at http://www.wormatlas.org/images/NeuronImageList.jpg.

### Soma and process morphology of amphid interneurons are conserved

We next examined the next processing layer of amphid sensory information in the worm, the first-layer amphid interneurons. In *C. elegans* there are four such amphid interneuron classes, each composed of a bilaterally symmetric pair of neurons (termed AIA, AIB, AIY, AIZ, for <u>A</u>mphid Interneuron <u>A</u>, <u>B</u>, <u>Y</u> and <u>Z</u>). All 4 neurons display highly distinctive features and an examination of the *P. pacificus* nervous system revealed a remarkable extent of conservation of these features, thereby easily identifying these neurons (the ease of identification prompted us to skip the tentative numerical naming scheme that we initially applied to sensory neurons, but rather give these *P. pacificus* neurons the same name as in *C. elegans* straight away). In both species, the AIY interneuron cell bodies are located in the same relative position in the ventral ganglion and their axons displays a characteristic ‘humped’ morphology (White et al., 1986). This ‘humped’ morphology corresponds to a large synaptic output onto AIZ, which forms a sheet-like cross-section immediately dorsal to AIY (Fig. 7B and 7D). The AIB neurons are identifiable by a characteristic switch of their process into two distinct neighborhoods of the anterior portion of the ventral ganglion, where its proximal axon is ventral and distal axon is more dorsal. The somas of the AIA interneurons occupy a stereotypic mediodorsal location and, like the AIY and AIZ neurons, form a dorsal midline gap junction. We rendered the complete structure of the amphid interneurons in 3D, further illustrating that cell body locations and axon projections are nearly identical in both species (Fig. 7F).

### Patterns of conservation and divergence in synaptic connectivity

Having identified the first-layer amphid interneurons in *P. pacificus,* we next explored the degree to which synaptic connectivity in the amphid circuit is conserved across species. We annotated all chemical synapses and gap junctions between the amphid sensory neurons, AUA, AIA, AIB, AIY, and AIZ, recording both the number of individual synapses as well as the number of serial section electron micrographs where ultrastructural synaptic anatomy was present as a proxy for anatomical connection strength. We identified 138 chemical connections (directed edges in a graph of connectivity) and 98 gap junction connections (undirected edges) in *P. pacificus*, compared to 73 chemical edges 96 gap junction edges in a recent re-evaluation of connectivity in *C. elegans* (Cook et al 2019, in press). As small synapses are more difficult to annotate reliably (Xu et al., 2013), we limited our analysis to only include connections that are ≥10 EM sections in strength in both species, which represents the strongest 50% of synaptic connections. Such thresholding also should minimize any potential concerns about comparing synaptic annotations between distinct datasets of different provenance.

Of these strong chemical synaptic connections, 32/53 connections were present in both species while 15 and 6 were specific to *P. pacificus* and *C. elegans*, respectively. Four of the strong gap junction connections were present in both species while 6 and 2 were specific to *P. pacificus* and *C. elegans*, respectively (Fig. 8A). To better contextualize similarities and differences in connectivity, we created a circuit diagram that shows a layered output from sensory neurons (triangles) onto interneurons (hexagons). We found that, on average, conserved edges are larger, more frequently made by amphid interneurons, and neurons whose structure is qualitatively most similar between species made more similar synaptic outputs (Fig. 8B). Examples include the synaptic output of AFD, whose output is almost exclusively onto AIY, or the patterns of interconnectivity of the amphid interneurons. In contrast, the amphid wing neurons (AWA, AWB, AWC) showed multiple connections present in only one species. For example, AWA makes strong synaptic connections to the AIB interneurons exclusively in *P. pacificus.* Similarly, the *P. pacificus* polymodal ASH neuron class also shows distinct patterns of synaptic connectivity: ASH makes *P. pacificus*-specific outputs to AIY and is innervated by AIA. While many of the *P. pacificus* synaptic contacts concern the connection of sensory neurons to first layer interneurons, many of the *C. elegans-*specific synaptic contacts concern the connections between sensory neurons. This may indicate an increase in cross-communication of sensory modalities in *C. elegans* as well as distinctions in which sensory outputs are processed in *P. pacificus*.

**Figure 8.**
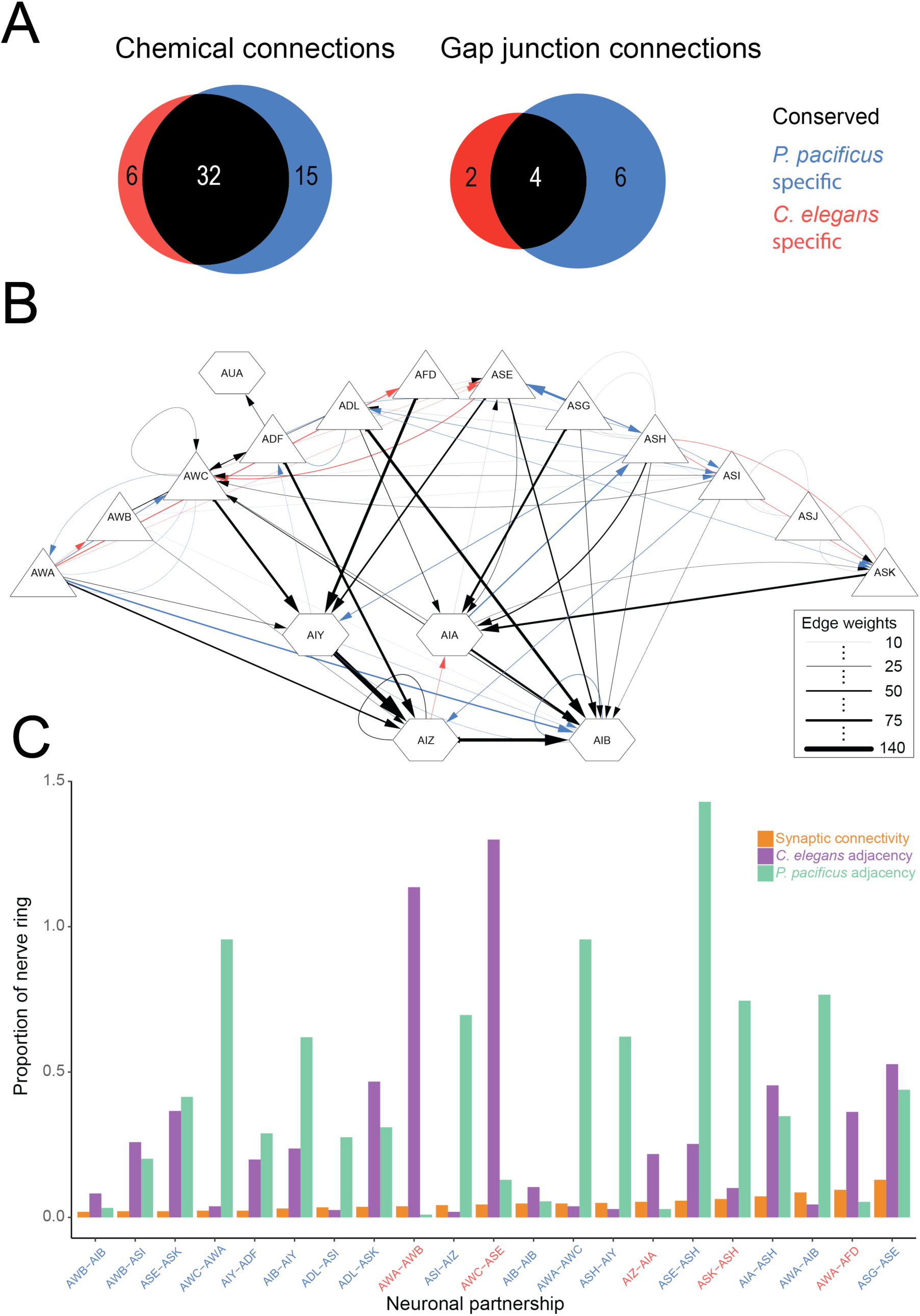
(A) Venn diagrams showing number of conserved (black), *C. elegans*-specific (red), and *P. pacificus*-specific (blue) chemical connections (directed edges) and gap junction connections (undirected edges). Threshold for edge weight was set to >= 10 serial EM sections, connectivity was determined between amphid sensory neurons, AUA, and amphid interneurons AIA, AIB, AIY, and AIZ. (B) Circuit diagram showing sensory neurons (triangles) connecting to interneurons (hexagons) by chemical synaptic connectivity (arrows) and gap junctions (lines). The weight of connection (number of serial section electron micrographs for all connections ≥10 sections). (C) Bar graph showing connectivity (orange), neuronal adjacency for *C. elegans* (purple), and neuronal adjacency for *P. pacificus* (green). Adjacency was determined computationally by comparing adjacent pixels of volumetrically traced neuron profiles. Proportion was determined by dividing the number of serial EM sections showing a synaptic connection or adjacency by the total number of sections comprising the nerve ring and ventral ganglion. Each partnership evaluated is a combination of two neurons who formed a species-specific connection shown in (B).

Synaptic connections present in only one species could be due to either differences in the structure of neuronal neighborhoods or synaptic specificity among adjacent processes. To distinguish between these possibilities, we computationally determined all axon-axon adjacencies of the amphid circuit in both species as previously described (Brittin et al., 2018). To compare across species, and to reduce section thickness and fixation artifacts, we calculated the proportion of adjacency (total number of EM sections where two processes are adjacent divided by the number of sections in the region of interest). Of the 29 connections we scored present only in one species, there are only 21 possible neuron-neuron adjacencies due to reciprocal connectivity. Of these 21 neuron-neuron pairs, we found that all make physical contact in both species. The amount of contact did, however, vary across a wide range in both species. Among species-specific neuronal partnerships, we found the percent of axonal-axonal contact ranged from 0.95% to 142% (median contact 26.7%) (Fig 8C). We evaluated whether the amount of neuron-neuron adjacency could predict the strength of a synaptic connection. We compared the correlation between synaptic connectivity and neuron-neuron adjacency and found a very weak, non-significant, correlation in both species (Spearman’s R = 0.2042, p=0.3747 in *P. pacificus;* Spearman’s R = 0.1248, p=0.5898 in *C. elegans*). Together, these results suggest that species-specific differences in connectivity are largely determined by synaptic partnership choice or recognition rather than large differences in axonal fasciculation and neuronal neighborhood changes.

### Similarity and divergences of molecular features

Moving beyond anatomy, we finally assessed the evolutionary divergence of molecular features of individual neurons. Divergences in molecular features between homologous neurons in different species can be expected to be manifested via the species-specific loss or gain of genes that control certain neuronal phenotypes or via the modifications of expression patterns of genes. There appears to be ample evidence for both. For example, several census taken for the number of GPCR-type sensory receptor-encoding genes indicated a much smaller number in *P. pacificus* compared to *C. elegans* (Dieterich et al., 2008; Krishnan et al., 2014). Similarly, there are species-specific gene losses and gain in the complement of specific subfamilies of taste receptor-type guanylyl cyclases, as discussed above. On the level of regulatory factors, we noted the absence of the *lsy-6* locus outside the *Rhabditiae*. In regard to genes that control structural features to neurons, we note that a gene, *oig-8*, that was recently identified as controlling the morphological elaborations of the winged cilia of the AWA/B/C neurons in *C. elegans* (Howell and Hobert, 2017), is also not present in the *P. pacificus* genome. This gene, *oig-8*, encodes a single transmembrane, immunoglobulin domain protein. To assess whether this gene loss is responsible for the lack of winged cilia in *P. pacificus,* we mis-expressed *Cel-oig-8* under the *Ppa-odr-7* promoter (described below) along with the *Ppa-odr-7p::rfp* marker but found no difference in the terminal dendritic cilia of RFP positive neurons that were co-injected with the *Ppa-odr-7p::Cel-oig-8* transgene compared to those that were injected only with the *Ppa-odr-7p::*RFP marker (SI Fig. 7). This result suggests the two *Ppa-odr-7*-expressing amphid neurons may lack other factors, such as terminal selector transcription factors, to realize the branching function of *Cel-oig-8* (Howell and Hobert, 2017).

To assess potential divergences in gene expression patterns, we considered three phylogenetically conserved genes that encode for regulatory and signaling factors. For the regulatory factors, we considered two transcription factors that are expressed exclusively in a single neuron class in *C. elegans*, *odr-*7, an orphan nuclear hormone receptor uniquely expressed in AWA (Sengupta et al., 1994) and *che-1,* uniquely expressed in the ASE neurons (Tursun et al., 2009). Both transcription factors control the differentiated state of the respective neuron class in *C. elegans* (Etchberger et al., 2007; Sengupta et al., 1994; Uchida, 2003) and 1-1 orthologs could be identified in *P. pacificus* by reciprocal best BLASTP hits as well as protein sequence phylogeny (SI Fig. 8A-B). We generated a reporter gene fusion for the *Ppa-che-1* locus and found that transgenic animals expressed the reporter in two neuron pairs in the head region of the worm (SI Fig. 9). The same expression can be observed by smFISH against the endogenous locus (SI Fig. 10). By cell positions, as well as by the lack of axon terminating pass the dorsal midline using fluorescence microscopy, we identified these neurons as the AS5(ASE) neurons and the AS6(ASG) neurons. However, we attempted but could not further confirm that the ASG axons terminate at the lateral midline. Expression in AS5(ASE) matches the expression observed in *C. elegans* and expression in AS6(ASG) is notable because the ASG neuron is also a water-soluble chemical taste receptor (Bargmann and Horvitz, 1991) that can compensate for loss of ASE function under hypoxic conditions (Pocock and Hobert, 2010). Since the AS5(ASE) neurons of *P. pacificus* may not be functionally lateralized and, therefore, may not be able to discriminate between cues that are sensed by ASEL and ASER in *C. elegans*, we are inclined to speculate that *che-1* may endow AS6(ASG) neurons of *P. pacificus* with chemosensory abilities that allow for the discrimination of ASE- and ASG-sensed water-soluble cues.

The expression pattern of the *odr-7* transcription factor seems to have diverged more substantially between *P. pacificus* and *C. elegans.* Based on cell body position and axonal projections, a *Ppa-odr-7p::rfp* reporter is expressed in the proposed AS7(AWC) and the ADF homolog AD9(ADF) neuron pair (Fig. 3 G-H, SI Fig. 11). This divergence of expression is particularly notable if one considers that *C. elegans odr-7* contributes to the specification of *C. elegans-*specific elaboration of AWA cilia (Howell and Hobert, 2017). In contrast, *P. pacificus* AD3(AWA) does not express *odr-7* and its cilia do not display the winged elaborations, as discussed above. Since, unlike with *che-1*, we have not confirmed endogenous *odr-7* expression with smFISH, it is conceivable that our reporter construct lacks relevant cis-regulatory elements.

In contrast to the reduction of *gcy* genes important for *C. elegans* ASE laterality, the *P. pacificus* genome contains almost all of the downstream G-protein subunit signaling proteins found to be encoded in the *C. elegans* genome (SI Fig. 8C). We therefore established a transgenic reporter strain using the promoter of *Ppa-odr-3*, a G-protein subunit homolog known to be expressed in a number of *C. elegans* sensory neurons, most strongly in AWC, weaker in AWB, and faintly in AWA, ADF, ASH neurons in *C. elegans* (Roayaie et al., 1998). We found consistent *Ppa-odr-3p::rfp* expression in AD3(AWA) and less robust expression in AS4(ASK)(Fig. 3C-D). Hence, like *odr-7* expression, the conservation of cell-type specific expression is limited to the amphid neurons but not to the proposed cellular homologs when soma position and axon projection patterns are also considered.

## Conclusions

The genomes of *C. elegans* and *P. pacificus* are remarkably distinct. Detailed recent analysis using phylotranscriptomics, Illumina, and single molecule sequencing revealed striking differences in gene content and genome organization to more accurately date their divergence to ~100 million year ago (Prabh et al., 2018; Rodelsperger, 2018; Rodelsperger et al., 2017). In light of this divergence, the extent of similarities of nervous system patterning is remarkable. Neuron number appears invariant, neuronal soma position is restricted, and perhaps most remarkable are the similarities in process outgrowth and relative process position. Relative neighborhoods of processes are retained, including a number of remarkably subtle aspects of process morphology, such as the highly unique and characteristic neighborhood change of the AIB processes or the humped axon morphology of AIY at a specific location. These findings strongly suggest the existence of constraints in patterning of the nervous system, particularly during the phase of axonal and dendritic outgrowth, i.e. relative placement of processes into specific neighborhoods. Such placement is a critical pre-requisite for proper synaptic targeting choice by constraining which possible targets any given neuron can innervate (White, 1985). In light of these constraints, it is enlightening to observe a number of striking differences in synaptic connectivity. These changes are likely to generate very distinct avenues of information flow, thereby altering behavior. Taken together, our results argue that alterations in synaptic connectivity, rather than initial patterning of neuronal fascicles are a key substrate of evolutionary change.

There are genomic differences between *P. pacificus* and *C. elegans* that display tantalizing correlates to some of the specific neuroanatomical diversities. The three main olfactory neurons of *C. elegans* display elaborated cilia morphology (winged cilia) where olfactory receptors are known to localize. The *C. elegans* genome encodes more than 1300 olfactory-type GPCRs (Robertson and Thomas, 2006; Troemel et al., 1995), many known to be co-expressed in AWA, AWB and AWC (Troemel et al., 1995; Vidal et al., 2018). In contrast, the *P. pacificus* genome contains many fewer olfactory-type GPCRs than *C. elegans* (Prabh et al., 2018), correlating with less morphological complexity of olfactory cilia. It is tempting to speculate that the expansion of the olfactory receptor repertoire in *C. elegans,* relative to *P. pacificus*, and the concomitant expansion of the morphological elaborations in *C. elegans* are functionally coordinated. Yet despite the smaller GPCR repertoire, *P. pacificus* exhibits an odor preference profile that has scant overlap with *C. elegans* (Cinkornpumin et al., 2014; Hong and Sommer, 2006), hence one priority for future studies is to identify the amphid neurons that express odor receptors in *P. pacificus*.

Other genome sequence changes reveal intriguing correlates to a specific synaptic alteration that we observed. Like many other neuron pairs, the AS5(ASE) neurons of *P. pacificus* are electrically coupled. In *C. elegans,* the left and right ASE neurons are not electrically coupled (even though the processes are in contact to each other), thereby allowing both sensory neurons to discriminate between distinct sensory cues sensed by the left and right ASE neurons (Ortiz et al., 2009; Suzuki et al.). Concomitant with the lack of electrical coupling of ASEL and ASER, a specific subset of ASEL and ASER-expressed rGC-type receptor proteins have expanded in the *C. elegans* genome (Ortiz et al., 2006), thereby expanding the spectrum of chemosensory cues that can be differentially sensed by ASEL vs. ASER. The genetic mechanisms to express these expanded set of *C. elegans-*specific receptor protein subfamilies in the left versus right ASE protein is triggered by the *lsy-6* miRNA, the most upstream regulator of ASEL/R asymmetry (Cochella and Hobert, 2012; Johnston and Hobert, 2003).. Remarkably, this miRNA is a *Caenorhabditis* genus-specific “invention”, i.e. it does not exist in *P. pacificus* (Ahmed et al., 2013). Future behavioral studies are required to conclusively demonstrate that our neuroanatomical studies have indeed revealed the cellular, genetic and anatomical means by which functional laterality arose in *C. elegans*.

## Materials and Methods

### Transmission electron microscopy

The TEM data used in the current study are derived from two sets of slightly different preparations of young adult *P. pacificus* strain PS312 hermaphrodites. All of the 3D reconstructions and most of the morphological details originate from two datasets of roughly 3000 serial TEM sections of 50 nm thickness each covering the anterior parts of two high-pressure-frozen and freeze-substituted adult hermaphrodites (specimen 107 and specimen 148), which were generated by Daniel Bumbarger. For a detailed methods description see Bumbarger and coworkers (Bumbarger et al., 2013), for even greater detail refer to protocol 9 of the *Pristionchus pacificus* protocols in the Methods for nematode species other than *C. elegans* section by Pires da Silva (2013) in WormBook (doi/10.1895/wormbook.1.7.1, http://www.wormbook.org). Alignment and manual segmentation was done in TrakEM2 (Cardona et al., 2012; Kremer et al., 1996). Segmentation and 3D reconstruction of the sensory head neurons were initially performed by Tahmineh Sarpolaki (2011) in the course of her Diploma thesis. Analysis of neuronal adjacencies was performed by using a modified python script written by Christopher Brittin (Brittin et al., 2018) Circuit diagrams were generated using Cytoscape (Shannon et al., 2003) and graphs were generated using the ggplot2 package for R (Wickham, 2016).

We additionally used four specimens, three transversely sectioned and one sagittally, which were prepared as follows: worms were placed in 100 µm deep specimen carriers half-filled with thick *E. coli* OP50 suspension, covered with the flat side of another carrier and high-pressure-frozen with a Bal-tec HPM-10 high-pressure freezer (Balzers, Liechtenstein). Freeze substitution was carried out in a freeze-substitution unit (Balzers FSU 010, Bal-Tec, Balzers, Liechtenstein) <u>accord</u>ing to the following protocol: fix in 2% OsO_4_, 0.5% UA, 0.5% GA in 97.5% acetone, 2.5% Methanol for 24 h at −90 °C, raise temperature to −60 °C in 3 h, hold for 6 h, raise to −40 °C in 2 h, hold for 12 h, keep on ice for 1 h, wash with 100% acetone, embed in Epon/acetone. Blocks were sectioned with an LKB 2128 Ultratome. Ultrathin sections were viewed in a Philips CM10 or in a Fei Tecnai G2 Spirit T12 transmission electron microscope, images were acquired on photo plates or with a Morada TEM CCD camera, respectively.

### Scanning electron microscopy

Clean specimens of adult *P. pacificus* strain PS312 Sommer *et al*. (1996) were fixed in 2.5% glutaraldehyde in PBS, post-fixed with 1% osmium tetroxide in PBS, dehydrated in a graded series of 30%, 50%, 70%, 95% and 100% ethanol, critical-point dried in liquid CO_2_ and sputter-coated with 10 nm Au/Pd. Inspection was carried out at 15 kV in a Hitachi S-800 field emission scanning electron microscope. We viewed ten processed *P. pacificus* PS312 hermaphrodite adults “face on” and did not notice any differences in the external structures of the mouth regions among them.

### Transgenic reporter strains

*P. pacificus* California PS312 and transgenic strains were raised on OP50 *E. coli* seeded NGM plates at 20°C. We used *P. pacificus* gene names (PPAxxxxx) and putative homology from www.wormbase.org based on the most recent *P. pacificus* Genome Assembly El_Paco (Rodelsperger et al., 2017) and *C. elegans* WS269. To confirm the assigned gene orthology, we looked for best reciprocal BLASTP hits between *P. pacificus* and *C. elegans* genomes, as well as 1-1 orthology in gene phylogeny trees. To resolve ambiguities, we also performed BLASTP search against AUGUSTUS gene predictions in the previous *P. pacificus* Genome Assembly Hybrid1 (www.pristionchus.org>Genome>Hybrid1, select track AUGUSTUS2013) and the amino acid sequences downloaded under “Sequences”. To make a *Ppa-odr-3* reporter plasmid, a ~1.7 kb long region upstream of the first ATG codon of *Ppa-odr-3* (PPA14189) (FP: GAGCGAGTGAAATGAGCTCAGTCC, RP: GGGTGATCGATACGAGGAGTGTTC) and the coding sequence of TurboRFP fused to the 3’ UTR of the ribosomal gene *Ppa-rpl-23* (Schlager et al., 2009) were cloned into the pUC19 plasmid using Golden Gate Assembly Mix (New England BioLabs, E1600S) following the manufacturer’s instructions. To make the *Ppa-daf-6* reporter, a 2.4 kb promoter sequence upstream of the start codon *Ppa-daf-6* (PPA15978) (FP: CTCGCCCGTGGATCATGTG, RP: TGCAAATCATTGATTGAATCATGG) was fused with *rfp* and *Venus* by fusion PCR (Hobert, 2002; Kieninger et al., 2016). Because the homology assignment for *odr-7* on Wormbase.org was *Ppa-nhr-66*, we looked for more likely orthology candidates using the AUGUSTUS genome assembly. The best BLASTP hit of *Cel-odr-7* was Contig1-aug1055.t1 (Contig1:2723982-2725788) and the *Ppa-odr-7* promoter was amplified and fused to *rfp* (FP: AACCAATGCATTGGCTTAGTTGGTTTCACTAATCACTACTG, RP: CCCTTGTCATTCAGATGAGCGAGCTGATCAAGGAG). To test if *Cel-oig-8* expression in the *Ppa-odr-7-*expressing neurons is sufficient to induce branching, we fused the 1.8 kb *Ppa-odr-7* promoter region to the genomic region of *Cel-oig-8* (FP: CCCTTGTCATTCAGATGAGCCTCCTTTCCAATATT, RP: TTACAGGGAGAAAGAGCATGTAG) and injected it with the *Ppa-odr-7p::rfp*. Each fusion junction site was verified by Sanger sequencing. For the construction of *Ppa-che-1p::rfp*, a 3.1 kb upstream fragment containing the first exon was amplified and fused with *rfp*. Although *Ppa-che-1*(PPA01143) was not the best hit when using the *Cel-che-1* as query, *Cel-che-1* was the best hit when using PPA01143 as query. The best hit homolog on Wormbase.org, *Ppa-blmp-1*(PPA04978), is predicted to encode for a much larger protein than both *che-1* and *Ppa-che-1*(PPA01143). Curiously, no RFP fluorescence was visible when the reporter did not include the endogenous first exon and intron fused to *rfp*. The expression pattern observed with this transgene recapitulated the pattern of expression of the endogenous *Ppa-che-1* determined by single-molecule fluorescence *in situ* hybridization (smFISH). The probe set used to stain for *Ppa-che-1* was obtained from Stellaris, Biosearch Technologies (Middlesex, UK) and targets the validated, full-length *Ppa-che-1* sequence. The smFISH staining was performed as previously described (Ji and van Oudenaarde, 2012).

To create complex arrays for transgenesis, wildtype PS312 genomic DNA, the PCR product or the plasmid carrying the target reporter, and the *Ppa-egl-20::rfp* reporter (co-injection marker expressed in the tail) were digested with the FastDigest PstI restriction enzyme (Thermo Fisher Scientific, FD0615) and then mixed at the final concentration of 60 ng/µl for genomic DNA and 10 ng/µl for each plasmid. Prepared mix was injected in the gonad rachis in hermaphrodites (Cinkornpumin and Hong, 2011; Schlager et al., 2009). The F_1_ progeny of injected animals were examined under a fluorescent dissecting microscope and animals that expressed the *Ppa-egl-20p::rfp* co-injection marker were isolated. All reporters were maintained as extrachromosomal arrays: *Ppa-daf-6p::rfp* (*tuEx231*), *Ppa-daf-6p::venus* (*tuEx250*), *Ppa-che-1p::rfp (lucEx367), Ppa-odr-3p::rfp* (*tuEx265*) and *Ppa-odr-7p*::*rfp* (*tuEx296* and *tuEx297*).

### DiI live staining

Lipophilic dyes DiI and DiO (Molecular Probes, V22889 and V22886) were used as neuronal tracers for a stereotypical subset of amphid neurons to facilitate cell identification either by relative cell position or overlap of red or green fluorescence in combination with transgenic reporters. DiI (red) and DiO (green) specifically stain five head amphid neurons in *C. elegans*, but DiO stains only two (ADL and ASJ) of the five (ASH, ADL, ASK, ADF, and ASJ) possible stereotypical amphid neurons in *P. pacificus*. Well-fed nematodes were washed once in M9 buffer and then incubated for 2 hours at ~23°C with 300 µl of fresh M9 containing 1:150 dilution of DiI or DiO (6.7 mM). The nematodes were subsequently washed twice in 800 µl of fresh M9 buffer and placed onto OP50 *E. coli* seeded NGM plates to let the worms crawl freely for ~30 minutes to remove excess dye.

### Software

For images of DiI-filling in Figure 3A-B, we used AutoDeblur and Autovisualize v9.3 (AutoQuant Imaging, Inc., New York) to reduce fluorescence background, and MetaMorph v6.2r5 (Universal Imaging Corp. Pennsylvania) to 3D reconstruct or stack images from multiple planes. Adobe Photoshop and Image J were used to process other images.

## Acknowledgements

We would like to thank Juergen Berger for assistance with scanning electron microscopy, Heinz Schwarz and Brigitte Sailer for assistance with transmission electron microscopy, Martin Voetsch for help with illustrations, Melissa Culhane, Suryesh Namdeo and Hanh Witte for technical assistance. R.L.H. is supported by the National Institutes of Health Award SC3GM105579 and was a past recipient of the Alexander von Humboldt Short Stay Award. S.J.C. is supported by National Institutes of Health Fellowship 5F32MH1154328. We acknowledge the support of NVIDIA Corporation with the donation of a Titan V GPU used for this research.

## Supplementary Information

SI Fig. 1: Receptor-type guanylyl cyclases in *P. pacificus and C. elegans*
SI Fig. 2: Video: Z-stack images of DiI stained J3 larva (ventral view).
SI Fig. 3: Video: 3D rendering of *Ppa-daf-6::RFP* J4 with confocal microscopy
SI Fig. 4: Posterior-anterior path of amphid dendrite entry into sheath glia
SI Fig. 5: Video: 3D rendering of amphid neurons
SI Fig. 6: URX cilium panel
SI Fig. 7: *Ppa-odr-7p::Cel-oig-8* mis-expression
SI Fig. 8: G-protein subunits in *P. pacificus* and *C. elegans*
SI Fig. 9: Single molecule FISH of *Ppa-che-1*
SI Fig. 10: Video: Z-stack images of *Ppa-odr-7p::rfp*
SI Fig. 11: Video: Z-stack images of *Ppa-che-1p::rfp*

**SI Fig. 1:**
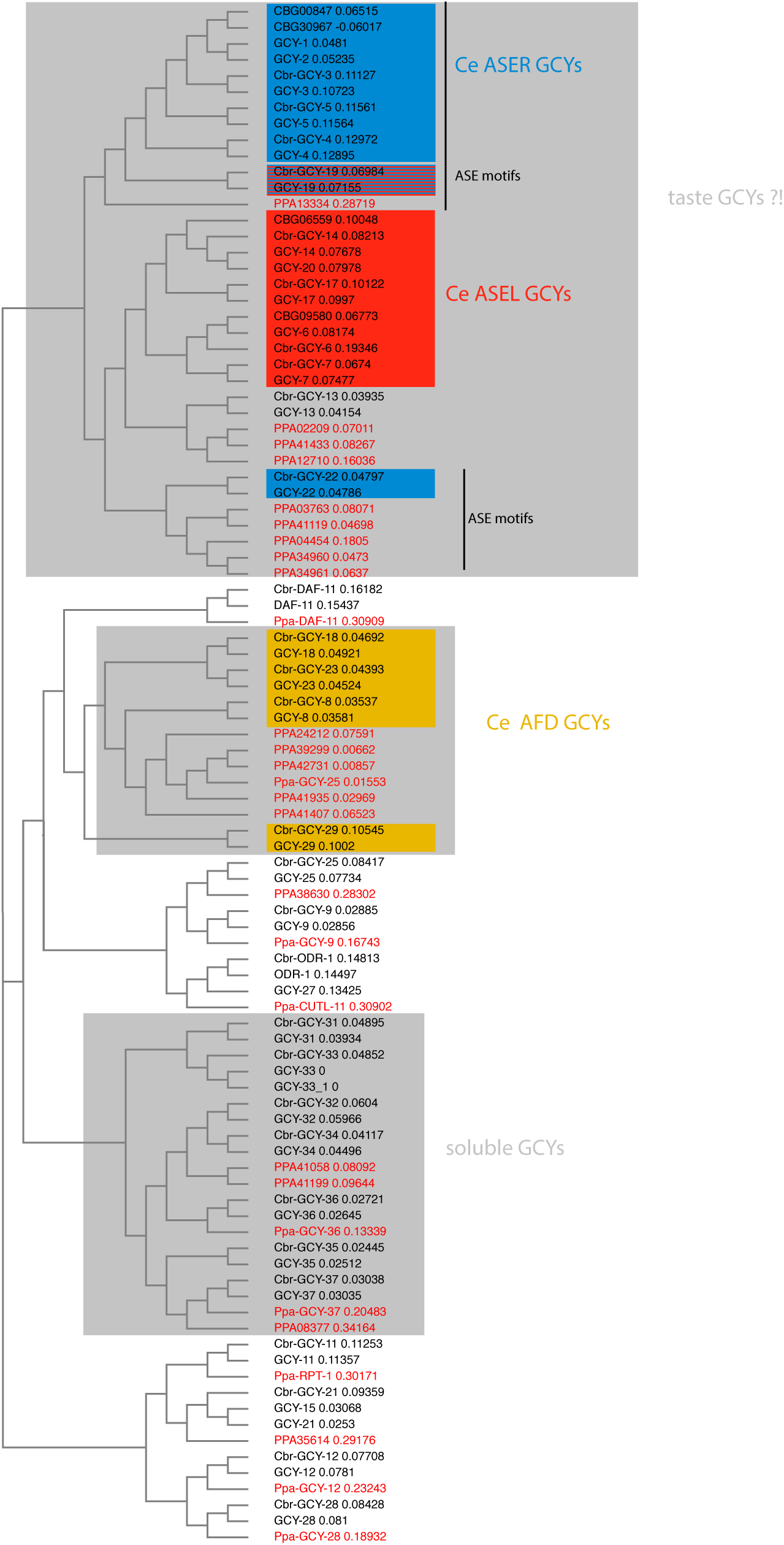
Receptor-type guanylyl cyclases in *P. pacificus* (red font) and *Caenorhabditis* species (black font).

SI Fig. 2: Video: Z-stack images of DiI stained J3 larva (ventral view). https://www.dropbox.com/s/6vdincoj29v014z/SI_Fig2_DiI_J3_11_ventral.m4v?dl=0

SI Fig. 3: Video: 3D rendering of *Ppa-daf-6::RFP* J4 with confocal microscopy https://www.dropbox.com/s/y5xg0vp9txwsi9l/SI_Fig3_Pdaf-6rfp_1J4.mp4?dl=0

**SI Fig. 4:**
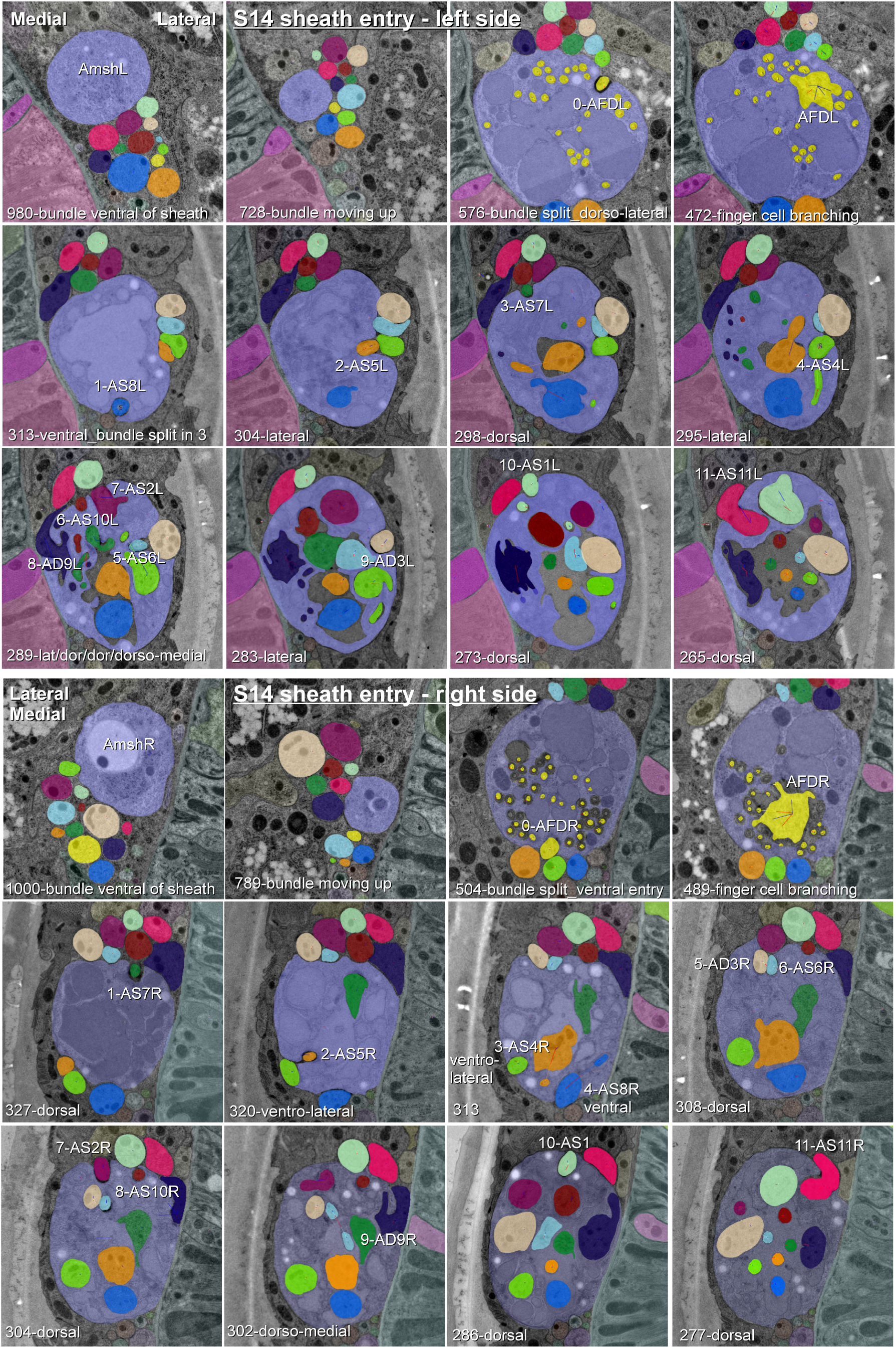
Posterior-anterior path of amphid dendrite entry into sheath glia. The entering dendrite is labeled according to *P. pacificus* nomenclature, section numbers from anterior and direction of entry is given at the bottom of each image.

SI Fig. 5: Video: 3D rendering of amphid neurons. https://www.dropbox.com/s/o7izf27y24p8uw8/SI_Fig5_amphid.epub?dl=0

**SI Fig. 6:**
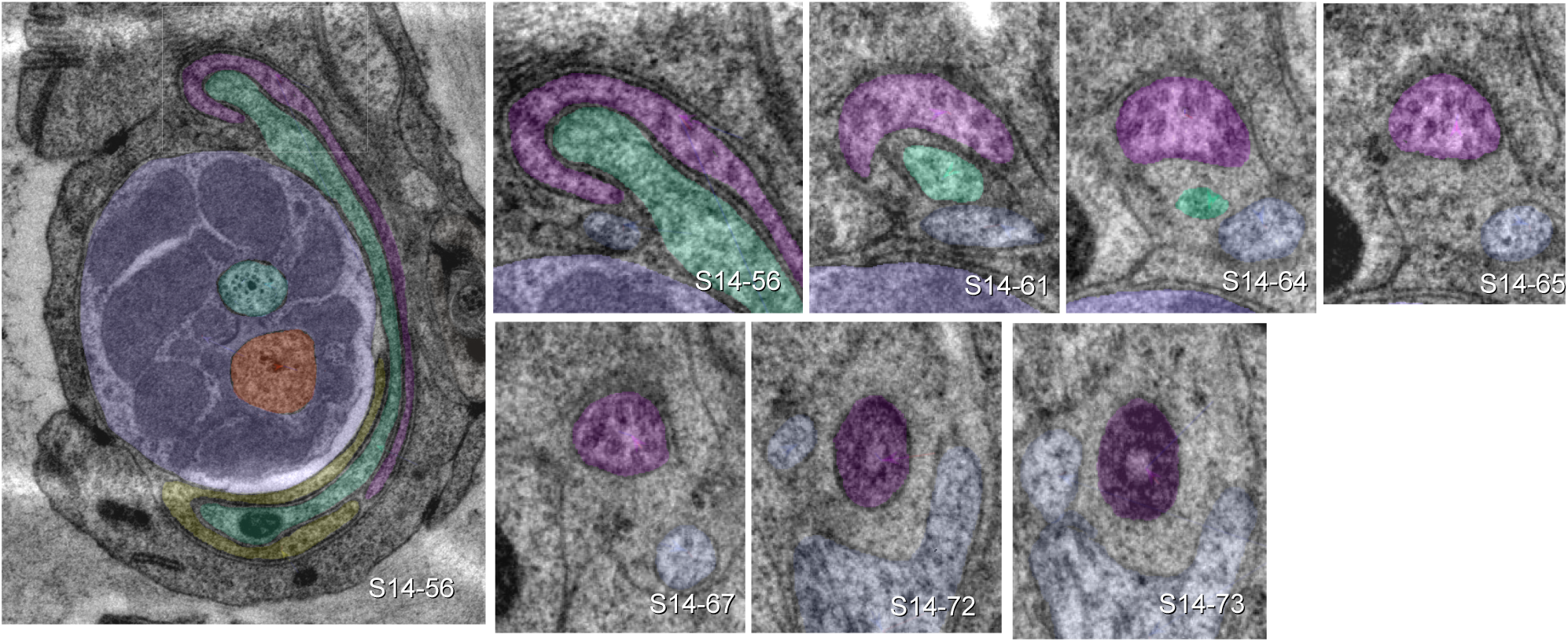
URX is ciliated. Large image shows left lateral IL sensillum of *P. pacificus* (specimen 107) with ILsoL (green), URXL (pink) and BAGL (yellow). Small images track the sheetlike URXL cilium from extended sheet-like tip (section 56) to transition zone (section 73). Sections are counted from the tip of the head. Dendritic structures can be compared to C. elegans at SlidableWorm http://www.wormatlas.org/SW/SW.php/ (slices 8-10).

**SI Fig. 7:**
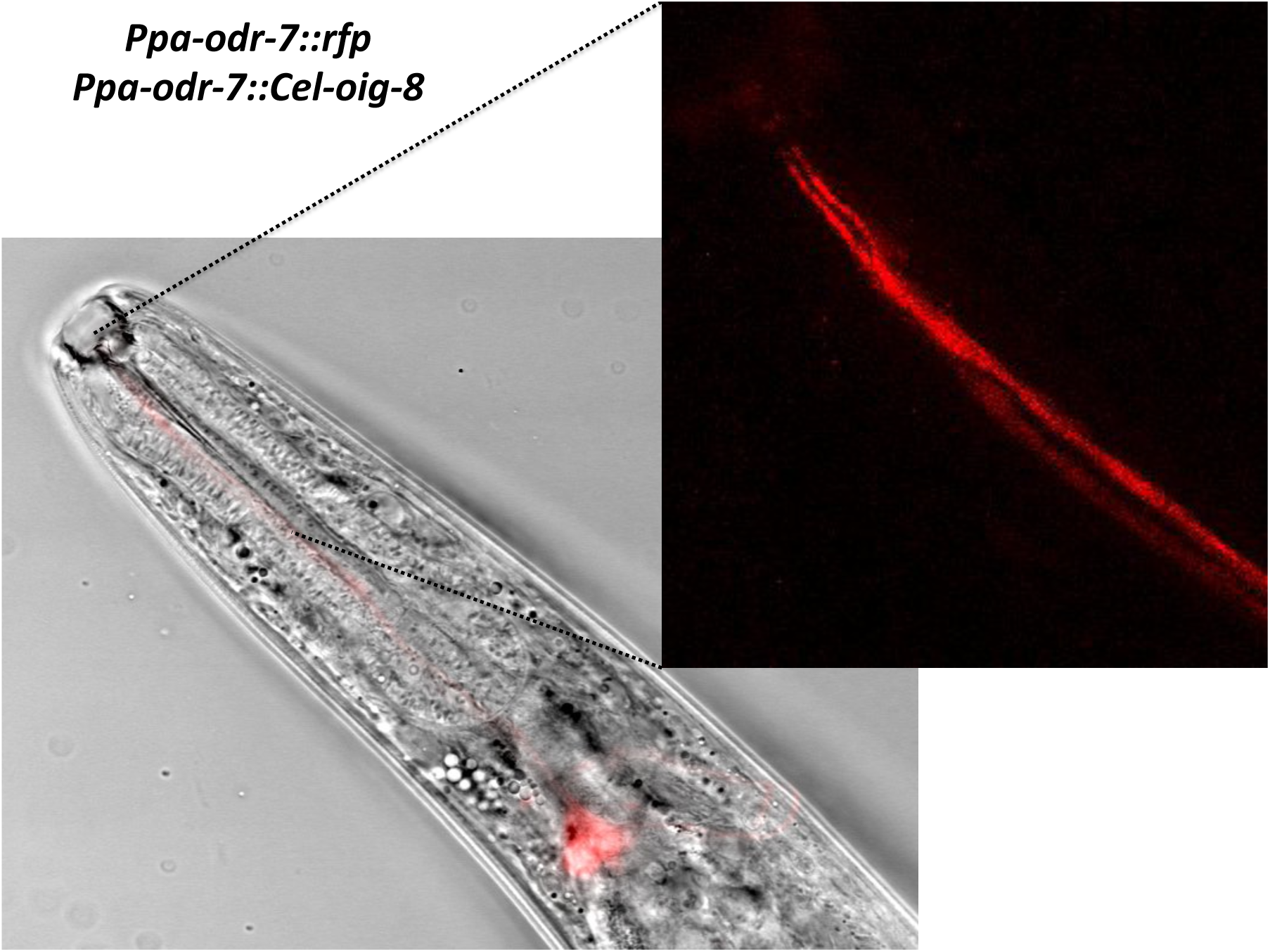
*Ppa-odr-7p::Cel-oig-8* mis-expression

**SI Fig. 8:**
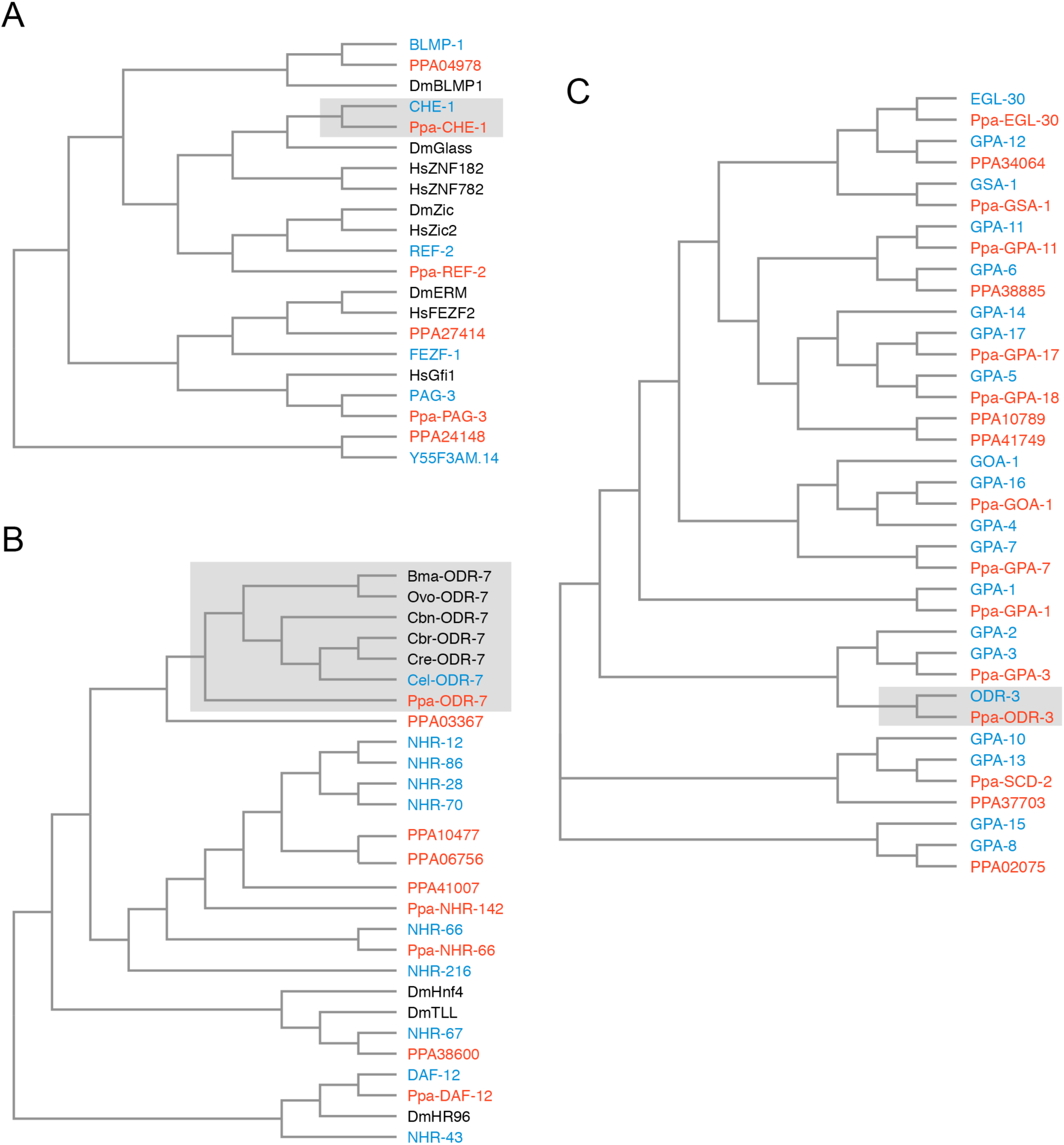
Orthology assignments for *che-1*, *odr-7* and *odr-3*. (A) Phylogenetic trees for the CHE-1 Zn finger transcription factor, (B) the ODR-7 nuclear hormone receptor, and (C) the ODR-3 G-alpha-protein subunits (shaded). The most closely related sequences were identified by reciprocal BLAST searches. Protein sequences for *P. pacificus* (red) are “Ppa-” (orthology same as Wormbase.org) or “PPA”, whereas the rest are *C. elegans* names (blue). Sequences from additional species (black) are *Drosophila melanogaster* (Dm), *Homo sapiens* (Hs), *Caenorhabditis briggsae* (Cbr), *Caenorhabditis remanei* (Cre), *Brugia malayi* (Bma), and *Onchocerca volvulus* (Ovo). For the nuclear hormone receptor tree building, only the DNA binding C4 domain was used. The trees were built by first aligning sequences with T-coffee (https://www.ebi.ac.uk/Tools/msa/tcoffee/), and the multiple sequence alignment was then fed into the Simply Phylogeny package in T-Coffee, with distance correction and exclusion of gaps.

SI Fig. 9: Video: Z-stack images of *Ppa-che-1p::rfp* https://www.dropbox.com/s/4kfx4jbdx15auuw/SI_Fig8_Pche1_DiO_J4_2stack_RH.m4v?dl=0

**SI Fig. 10:**
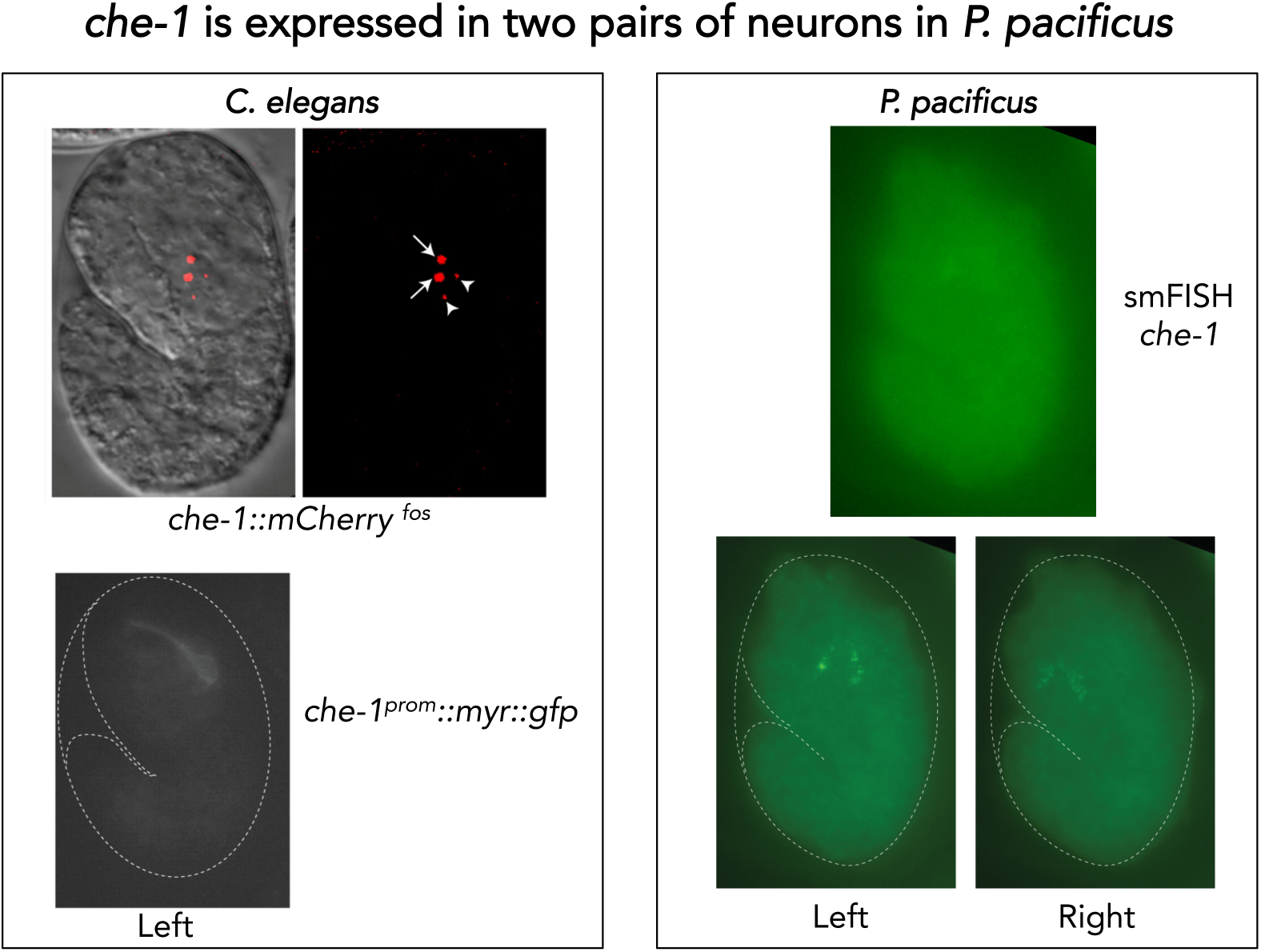
Single molecule FISH of *Ppa-che-1*. (Left) Representative images of *C. elegans* embryos carrying two different *Cel-che-1* reporters: a translational fusion of CHE-1 to mCherry in the context of a recombineered fosmid construct shows the expected nuclear localization in a pair of neurons identified as the ASE neurons (arrows) as well as two apoptotic bodies (arrow heads) corresponding to the dying ASE-sister cells. A 1.3 Kb fragment upstream of the *Cel-che-1* coding sequence also drives expression in the ASE neurons, in this case visualized with a myristoylated GFP reporter (Right) A representative *P. pacificus* embryo stained with the smFISH probe set against *Ppa-che-1*. Individual planes focused on the left or right sides of the embryo show two neurons on each side. The Z-stack series from which these images were extracted can be obtained at: link

SI Fig. 11: Video: Z-stack images of *Ppa-odr-7p::rfp* https://www.dropbox.com/s/6q6alebl1s2zqi3/SI_Fig10_Podr7RFP_Zstk.mov?dl=0

## References

Ahmed, R., Chang, Z., Younis, A. E., Langnick, C., Li, N., Chen, W., Brattig, N. and Dieterich, C. (2013). Conserved miRNAs Are Candidate Post-Transcriptional Regulators of Developmental Arrest in Free-Living and Parasitic Nematodes. Genome Biology and Evolution 5, 1246–1260.

Albeg, A., Smith, C. J., Chatzigeorgiou, M., Feitelson, D. G., Hall, D. H., Schafer, W. R., Miller, D. M. and Treinin, M. (2011). C. elegans multi-dendritic sensory neurons: Morphology and function. Molecular and Cellular Neuroscience 46, 308–317.

Albert, P. S. and Riddle, D. L. (1983). Developmental Alterations in Sensory Neuroanatomy of the Caenorhabditis-Elegans Dauer Larva. Journal of Comparative Neurology 219, 461–481.

Ashton, F. T., Bhopale, V. M., Fine, A. E. and Schad, G. A. (1995). Sensory Neuroanatomy of a Skin-Penetrating Nematode Parasite - Strongyloides stercoralis. I. Amphidial Neurons. Journal of Comparative Neurology 357, 281–295.

Bacaj, T., Tevlin, M., Lu, Y. and Shaham, S. (2008). Glia Are Essential for Sensory Organ Function in C. elegans. Science’s STKE 322, 744.

Bargmann, C. I. (2006). Chemosensation in C. elegans. WormBook 1–29.

Bargmann, C. I. and Horvitz, H. R. (1991). Chemosensory neurons with overlapping functions direct chemotaxis to multiple chemicals in C. elegans. Neuron 7, 729–742.

Bargmann, C. I., Hartwieg, E. and Horvitz, H. R. (1993). Odorant-selective genes and neurons mediate olfaction in C. elegans. Cell 74, 515–527.

Barrios, A., Ghosh, R., Fang, C., Emmons, S. W. and Barr, M. M. (2012). PDF-1 neuropeptide signaling modulates a neural circuit for mate-searching behavior in C. elegans. Nat Neurosci 15, 1675–1682.

Bento, G., Ogawa, A. and Sommer, R. J. (2010). Co-option of the hormone-signalling module dafachronic acid-DAF-12 in nematode evolution. Nature 466, 494–497.

Brittin, C. A., Cook, S. J., Hall, D. H., Emmons, S. W. and Cohen, N. (2018). Volumetric reconstruction of main *Caenorhabditis elegans* neuropil at two different time points. bioRxiv 485771.

Bumbarger, D. J., Crum, J., Ellisman, M. and Baldwin, J. G. (2007). Three-dimensional reconstruction of epidermal and sensory organs in the nematode Acrobeles complexus. Journal of Nematology 39, 87–87.

Bumbarger, D. J., Riebesell, M., Rodelsperger, C. and Sommer, R. J. (2013). System-wide Rewiring Underlies Behavioral Differences in Predatory and Bacterial-Feeding Nematodes. Cell 152, 109–119.

Bumbarger, D. J., Wijeratne, S., Carter, C., Crum, J., Ellisman, M. H. and Baldwin, J. G. (2009a). Three-dimensional reconstruction of the amphid sensilla in the microbial feeding nematode, Acrobeles complexus (Nematoda: Rhabditida). J Comp Neurol 512, 271–281.

Bumbarger, D. J., Wijeratne, S., Carter, C., Crum, J., Ellisman, M. H. and Baldwin, J. G. (2009b). Three-dimensional fine structural reconstruction of the nose sensory structures of Acrobeles complexus compared to Caenorhabditis elegans (Nematoda: Rhabditida). J Morphol 268, 649–663.

Cardona, A., Saalfeld, S., Schindelin, J., Arganda-Carreras, I., Preibisch, S., Longair, M., Tomancak, P., Hartenstein, V. and Douglas, R. J. (2012). TrakEM2 Software for Neural Circuit Reconstruction. PLoS ONE 7, e38011.

Carrillo, M. A., Guillermin, M. L., Rengarajan, S., Okubo, R. P. and Hallem, E. A. (2013). O2-Sensing Neurons Control CO2 Response in C. elegans. Journal of Neuroscience 33, 9675–9683.

Chang, A. J., Chronis, N., Karow, D. S., Marletta, M. A. and Bargmann, C. I. (2006). A distributed chemosensory circuit for oxygen preference in C. elegans. Plos Biology 4, 1588–1602.

Chatzigeorgiou, M. and Schafer, W. R. (2011). Lateral Facilitation between Primary Mechanosensory Neurons Controls Nose Touch Perception in C. elegans. Neuron 70, 299–309.

Cinkornpumin, J. K. and Hong, R. L. (2011). RNAi mediated gene knockdown and transgenesis by microinjection in the necromenic Nematode *Pristionchus pacificus*. J Vis Exp e3270.

Cinkornpumin, J. K., Wisidagama, D. R., Rapoport, V., Go, J. L., Dieterich, C., Wang, X., Sommer, R. J. and Hong, R. L. (2014). A host beetle pheromone regulates development and behavior in the nematode *Pristionchus pacificus*. eLife 3.

Coates, J. C. and de Bono, M. (2002). Antagonistic pathways in neurons exposed to body fluid regulate social feeding in *Caenorhabditis elegans*. Nature 419, 925–929.

Cochella, L. and Hobert, O. (2012). Embryonic Priming of a miRNA Locus Predetermines Postmitotic NeuronalLeft/Right Asymmetry in C. elegans. Cell 151, 1229–1242.

Dieterich, C., Clifton, S. W., Schuster, L. N., Chinwalla, A., Delehaunty, K., Dinkelacker, I., Fulton, L., Fulton, R., Godfrey, J., Minx, P., et al. (2008). The Pristionchus pacificus genome provides a unique perspective on nematode lifestyle and parasitism. Nat Genet 40, 1193–1198.

Doroquez, D. B., Berciu, C., Anderson, J. R., Sengupta, P. and Nicastro, D. (2014). A high-resolution morphological and ultrastructural map of anterior sensory cilia and glia in *Caenorhabditis elegans*. eLife 3.

Etchberger, J. F., Lorch, A., Sleumer, M. C., Zapf, R., Jones, S. J., Marra, M. A., Holt, R. A., Moerman, D. G. and Hobert, O. (2007). The molecular signature and cis-regulatory architecture of a C. elegans gustatory neuron. Genes Dev 21, 1653–1674.

Gray, J. M., Karow, D. S., Lu, H., Chang, A. J., Chang, J. S., Ellis, R. E., Marletta, M. A. and Bargmann, C. I. (2004). Oxygen sensation and social feeding mediated by a C. elegans guanylate cyclase homologue. Nature 430, 317–322.

Hallem, E. A. and Sternberg, P. W. (2008). Acute carbon dioxide avoidance in Caenorhabditis elegans. Proc. Natl. Acad. Sci. U.S.A. 105, 8038–8043.

Hallem, E. A., Spencer, W. C., McWhirter, R. D., Zeller, G., Henz, S. R., R H atsch, G., Miller, D. M., Horvitz, H. R., Sternberg, P. W. and Ringstad, N. (2011). Receptor-type guanylate cyclase is required for carbon dioxide sensation by Caenorhabditis elegans. Proc. Natl. Acad. Sci. U.S.A. 108, 254–259.

Han, Z., Boas, S. and Schroeder, N. E. (2016). Unexpected Variation in Neuroanatomy among Diverse Nematode Species. Front. Neuroanat. 9, 296.

Hedgecock, E. M., Culotti, J. G., Thomson, J. N. and Perkins, L. A. (1985). Axonal guidance mutants of Caenorhabditis elegans identified by filling sensory neurons with fluorescein dyes. Dev Biol 111, 158–170.

Heiman, M. G. and Shaham, S. (2009). DEX-1 and DYF-7 Establish Sensory Dendrite Length by Anchoring Dendritic Tips during Cell Migration. Cell.

Hobert, O. (2002). PCR Fusion-Based approach to create reporter gene constructs for Expression Analysis in Transgenic C. elegans. Biotechniques 32, 728–730.

Hong, R. L. and Sommer, R. J. (2006). Chemoattraction in Pristionchus nematodes and implications for insect recognition. Curr. Biol. 16, 2359–2365.

Howell, K. and Hobert, O. (2017). Morphological Diversity of C. elegans Sensory Cilia Instructed by the Differential Expression of an Immunoglobulin Domain Protein. Curr. Biol. 27, 1782–1790.e5.

Ji, N. and van Oudenaarde, A. (2012). Single molecule fluorescent *in situ* hybridization (smFISH) of *C. elegans* worms and embryos. WormBook, *ed.* The C. elegans Research Community, Wormbook.

Johnston, R. J. and Hobert, O. (2003). A microRNA controlling left/right neuronal asymmetry in Caenorhabditis elegans. Nature 426, 845–849.

Kieninger, M. R., Ivers, N. A., Rodelsperger, C., Markov, G. V., Sommer, R. J. and Ragsdale, E. J. (2016). The Nuclear Hormone Receptor NHR-40 Acts Downstream of the Sulfatase EUD-1 as Part of a Developmental Plasticity Switch in Pristionchus. Curr. Biol. 26, 2174–2179.

Koneru, S. L., Salinas, H., Flores, G. E. and Hong, R. L. (2016). The bacterial community of entomophilic nematodes and host beetles. Mol Ecol 25, 2312–2324.

Kremer, J. R., Mastronarde, D. N. and McIntosh, J. R. (1996). Computer Visualization of Three-Dimensional Image Data Using IMOD. Journal of Structural Biology 116, 71–76.

Krishnan, A., Almén, M. S., Fredriksson, R. and Schiöth, H. B. (2014). Insights into the Origin of Nematode Chemosensory GPCRs: Putative Orthologs of the Srw Family Are Found across Several Phyla of Protostomes. PLoS ONE 9, e93048.

Li, J., Zhu, X. D., Ashton, F. T., Gamble, H. R. and Schad, G. A. (2001). Sensory neuroanatomy of a passively ingested nematode parasite, Haemonchus contortus: Amphidial neurons of the third-stage larva. Journal of Parasitology 87, 65–72.

Liu, L., MacKenzie, K. R., Putluri, N., Maletić-Savatić, M. and Bellen, H. J. (2017). The Glia-Neuron Lactate Shuttle and Elevated ROS Promote Lipid Synthesis in Neurons and Lipid Droplet Accumulation in Glia via APOE/D. Cell Metab 26, 719–737.e6.

Liu, L., Zhang, K., Sandoval, H., Yamamoto, S., Jaiswal, M., Sanz, E., Li, Z., Hui, J., Graham, B. H., Quintana, A., et al. (2015). Glial Lipid Droplets and ROS Induced by Mitochondrial DefectsPromote Neurodegeneration. Cell 160, 177–190.

Liu, Z., Kariya, M. J., Chute, C. D., Pribadi, A. K., Leinwand, S. G., Tong, A., Curran, K. P., Bose, N., Schroeder, F. C., Srinivasan, J., et al. (2018). Predator-secreted sulfolipids induce defensiveresponses in C. elegans. Nat Commun 1–13.

Nei, M. and Glazko, G. (2001). Estimation of divergence times from multiprotein sequences for a few mammalian species and several distantly related organisms. Proceedings of the National Academy of Sciences 98, 1–2497–2502.

Oikonomou, G., Perens, E. A., Lu, Y., Watanabe, S., Jorgensen, E. M. and Shaham, S. (2011). Opposing activities of LIT-1/NLK and DAF-6/patched-related direct sensory compartment morphogenesis in C. elegans. PLoS Biol 9, e1001121.

Ortiz, C. O., Etchberger, J. F., Posy, S. L., Frokjaer-Jensen, C., Lockery, S., Honig, B. and Hobert, O. (2006). Searching for neuronal left/right asymmetry: genomewide analysis of nematode receptor-type guanylyl cyclases. Genetics 173, 131–149.

Ortiz, C. O., Faumont, S., Takayama, J., Ahmed, H. K., Goldsmith, A. D., Pocock, R., McCormick, K. E., Kunimoto, H., Iino, Y., Lockery, S., et al. (2009). Lateralized Gustatory Behavior of C. elegans Is Controlled by Specific Receptor-Type Guanylyl Cyclases. Current Biology 19, 996–1004.

Peckol, E. L., Troemel, E. R. and Bargmann, C. I. (2001). Sensory experience and sensory activity regulate chemosensory receptor gene expression in Caenorhabditis elegans. Proc. Natl. Acad. Sci. U.S.A. 98, 11032–11038.

Penkov, S., Ogawa, A., Schmidt, U., Tate, D., Zagoriy, V., Boland, S., Gruner, M., Vorkel, D., Verbavatz, J.-M., Sommer, R. J., et al. (2014). a wax ester promotes collective host finding in the nematode Pristionchus pacificus. Nature Chemical Biology 1–7.

Perens, E. A. and Shaham, S. (2005). C-elegans daf-6 encodes a patched-related protein required for lumen formation. Dev Cell 8, 893–906.

Perkins, L. A., Hedgecock, E. M., Thomson, J. N. and Culotti, J. G. (1986). Mutant sensory cilia in the nematode Caenorhabditis elegans. Dev Biol 117, 456–487.

Pocock, R. and Hobert, O. (2010). Hypoxia activates a latent circuit for processing gustatory information in C. elegans. Nat Neurosci 13, 610–614.

Prabh, N., Roeseler, W., Witte, H., Eberhardt, G., Sommer, R. J. and Rodelsperger, C. (2018). Deep taxon sampling reveals the evolutionary dynamics of novel gene families in Pristionchusnematodes. Genome Res.

Procko, C., Lu, Y. and Shaham, S. (2011). Glia delimit shape changes of sensory neuron receptive endings in C. elegans. Development 138, 1371–1381.

Rabinowitch, I., Chatzigeorgiou, M., Zhao, B., Treinin, M. and Schafer, W. R. (2014). Rewiring neural circuits by the insertion of ectopic electrical synapses in transgenic C. elegans. Nat Commun 5, 1–6.

Ragsdale, E. J. (2015). Pristionchus pacificus. (ed. Sommer, R. J. Brill Academic Publishers.

Ragsdale, E. J., Ngo, P. T., Crum, J., Ellisman, M. H. and Baldwin, J. G. (2009). Comparative, three-dimensional anterior sensory reconstruction of *Aphelenchus avenae*. The Journal of comparative neurology 517, 616–632.

Roayaie, K., Crump, J. G., Sagasti, A. and Bargmann, C. I. (1998). The G alpha protein ODR-3 mediates olfactory and nociceptive function and controls cilium morphogenesis in C. elegans olfactory neurons. Neuron 20, 55–67.

Robertson, H. M. and Thomas, J. H. (2006). The putative chemoreceptor families of C. elegans. WormBook, *ed.* The C. elegans Research Community, Wormbook.

Rodelsperger, C. (2018). Comparative Genomics of Gene Loss and Gain in Caenorhabditis and Other Nematodes. In Comparative Genomics: Methods and Protocols (eds. Setubal, J. C., Stoye, J., and Stadler, P. F., pp. 419–432. New York, NY: Springer New York.

Rodelsperger, C., Meyer, J. M., Prabh, N., Lanz, C., Bemm, F. and Sommer, R. J. (2017). Single-Molecule Sequencing Reveals the Chromosome-Scale Genomic Architecture of the Nematode Model Organism Pristionchus pacificus. CellReports 21, 834–844.

Rota-Stabelli, O., Daley, A. C. and Pisani, D. (2013). Molecular Timetrees Reveal a Cambrian Colonization of Land and a New Scenario for Ecdysozoan Evolution. Current Biology 23, 392–398.

Schafer, W. (2016). Nematode nervous systems. Current Biology 26, R955–R959.

Schlager, B., Wang, X., Braach, G. and Sommer, R. J. (2009). Molecular cloning of a dominant roller mutant and establishment of DNA-mediated transformation in the nematode Pristionchus pacificus. Genesis 47, 300–304.

Schmidt-Rhaesa, A. (2007). The Evolution of Organ Systems. Oxford University Press.

Sengupta, P., Colbert, H. A. and Bargmann, C. I. (1994). The C. elegans gene odr-7 encodes an olfactory-specific member of the nuclear receptor superfamily. Cell 79, 971–980.

Shannon, P., Markiel, A., Ozier, O., Baliga, N. S., Wang, J. T., Ramage, D., Amin, N., Schwikowski, B. and Ideker, T. (2003). Cytoscape: A Software Environment for Integrated Models of Biomolecular Interaction Networks. Genome Research 13, 2498–2504.

Srinivasan, J., Durak, O. and Sternberg, P. W. (2008). Evolution of a polymodal sensory response network. BMC Biol 6, 52.

Starich, T. A., Herman, R. K., Kari, C. K., Yeh, W. H., Schackwitz, W. S., Schuyler, M. W., Collet, J., Thomas, J. H. and Riddle, D. L. (1995). Mutations affecting the chemosensory neurons of Caenorhabditis elegans. Genetics 139, 171–188.

Suzuki, H., Thiele, T. R., Faumont, S., Ezcurra, M., Lockery, S. R. and Schafer, W. R. Functional asymmetry in Caenorhabditis elegans taste neurons and its computational role in chemotaxis. Nature 454, 114 EP –.

Troemel, E. R., Chou, J. H., Dwyer, N. D., Colbert, H. A. and Bargmann, C. I. (1995). Divergent seven transmembrane receptors are candidate chemosensory receptors in C. elegans. Cell 83, 207–218.

Tursun, B., Cochella, L., Carrera, I. and Hobert, O. (2009). A Toolkit and Robust Pipeline for the Generation of Fosmid-Based Reporter Genes in C. elegans. PLoS ONE 4, e4625.

Uchida, O. (2003). The C. elegans che-1 gene encodes a zinc finger transcription factor required for specification of the ASE chemosensory neurons. Development 130, 1215–1224.

Vidal, B., Aghayeva, U., Sun, H., Wang, C., Glenwinkel, L., Bayer, E. A. and Hobert, O. (2018). An atlas of Caenorhabditis elegans chemoreceptor expression. PLoS Biol 16, e2004218.

Ward, S., Thomson, N., White, J. G. and Brenner, S. (1975). Electron microscopical reconstruction of the anterior sensory anatomy of the nematode Caenorhabditis elegans. J Comp Neurol 160, 313–337.

Ware, R. W., Clark, D., Crossland, K. and Russell, R. L. (1975). The nerve ring of the nematode Caenorhabditis elegans: sensory input and motor output. J Comp Neurol 162.

Werner, M. S., Sieriebriennikov, B., Prabh, N., Loschko, T., Lanz, C. and Sommer, R. J. (2018). Young genes have distinct gene structure, epigenetic profiles, and transcriptional regulation. Genome Res 28, 1675–1687.

White, J. G. (1985). Neuronal connectivity in Caenorhabditis elegans. Trends Neurosci 8, 277–283.

White, J. G., Southgate, E., Thomson, J. N. and Brenner, S. (1986). The Structure of the Nervous-System of the Nematode Caenorhabditis-Elegans. Philosophical Transactions of the Royal Society of London Series B-Biological Sciences 314, 1–340.

Wickham, H. (2016). ggplot2: elegant graphics for data analysis (Use R!). 2nd ed. Springer.

Xu, M., Jarrell, T. A., Wang, Y., Cook, S. J., Hall, D. H. and Emmons, S. W. (2013). Computer Assisted Assembly of Connectomes from Electron Micrographs: Application to Caenorhabditis elegans. PLoS ONE 8, e54050.

Yu, S., Avery, L., Baude, E. and Garbers, D. L. (1997). Guanylyl cyclase expression in specific sensory neurons: A new family of chemosensory receptors. Proceedings of the National Academy of Sciences 94, 3384–3387.

